# CDKs-mediated phosphorylation of PNKP is required for end-processing of single-strand DNA gaps on Okazaki Fragments and genome stability

**DOI:** 10.1101/2021.07.29.452278

**Authors:** Kaima Tsukada, Tomoko Miyake, Rikiya Imamura, Kotaro Saikawa, Mizuki Saito, Naoya Kase, Masamichi Ishiai, Yoshihisa Matsumoto, Mikio Shimada

**Affiliations:** Laboratory for Zero-Carbon Energy, Institute of Innovative Research, Tokyo Institute of Technology, 2-12-1 Oookayama, Meguro-ku, Tokyo, 152-8550; National Cancer Center Research Institute, 5-1-1 Tsukiji, Chuo-ku, Tokyo, 104-0045, Japan

## Abstract

Polynucleotide kinase phosphatase (PNKP) has enzymatic activities as 3′ -phosphatase and 5′ - kinase of DNA ends to promote DNA ligation and repair. Here, we show that cyclin-dependent kinases (CDKs) regulate the phosphorylation of threonine 118 (T118) in PNKP. This phosphorylation allows recruitment to the gapped DNA structure found in single-strand DNA nicks and/or gaps between Okazaki fragments (OFs) during DNA replication. T118A (alanine)-substituted PNKP-expressing cells exhibited accumulation of single-strand DNA gaps in S phase and accelerated replication fork progression. Furthermore, PNKP is involved in poly (ADP-ribose) polymerase 1 (PARP1)-dependent replication gap filling as a backup pathway in the absence of OFs ligation. Altogether, our data suggest that CDK-mediated PNKP phosphorylation at T118 is important for its recruitment to single-strand DNA gaps to proceed with OFs ligation and its backup errors via the gap-filling pathway to maintain genome stability.

## Introduction

Genomic DNA is threatened by intrinsic factors such as oxidative stress derived from mitochondria-dependent energy metabolism and extrinsic factors such as ionizing radiation, ultraviolet radiation, and chemical compounds. These factors generate various types of DNA damage, such as base damage, DNA single-strand breaks (SSBs), and double-strand breaks (DSBs). Because DNA lesions cause chromosomal aberrations, leading to aneuploidy and tumorigenesis, DNA repair machinery has evolved in living organisms. Polynucleotide kinase phosphatase (PNKP) is a key enzyme with a dual role that bears 3′ -phosphatase and 5′ -kinase activity ^1, 2, 3^. Human PNKP consists of 521 amino acids, including the forkhead-associated (FHA) domain (amino acid residues 1–110) in the amino-terminal region, and phosphatase (146–337) and kinase (341–516) domains in the carboxy-terminal region, which are connected by a linker region (111–145). PNKP is recruited to SSBs and DSBs sites depending on its interactions via the FHA domain of PNKP with XRCC1 and XRCC4, respectively, and is involved in base excision repair (BER), SSB repair, and non-homologous end joining (NHEJ) for DSB repair ^4, 5, 6, 7, 8^. The PNKP linker region includes a nuclear localization signal and phosphorylation sites ^7^. Phosphorylation of PNKP at serine 114 by ataxia telangiectasia mutated (ATM) is required for protein stability and efficient DNA repair for cellular survival ^9, 10^. Mutations in PNKP are associated with the human inherited disease microcephaly and seizures (MCSZ), a neurodevelopmental disease ^11^, ataxia oculomotor apraxia 4 (AOA4) ^12^, and Charcot-Marie-Tooth disease (CMT2B2), a neurodegenerative disease ^13^. These mutations are mostly located in the phosphatase or kinase domains and attenuate the phosphatase and kinase activities ^14, 15, 16^.

The BER and SSBs repair intermediates form gapped DNA structures, which are the main targets of PNKP. These DNA gaps are also found in Okazaki fragments (OFs) during DNA replication. DNA replication integrity is essential for ensuring genomic stability and accurate cell proliferation ^17, 18^. DNA replication is initiated at the origin of replication in the leading strand and at short RNA-primed DNA fragments, known as OFs, in the lagging strand ^19, 20^. Recent reports suggest that single-strand DNA gap-filling machinery is involved in OFs maturation ^21^. Poly (ADP-ribose) polymerase 1/2 (PARP1/2) is involved in the repair of SSBs, DSBs, and multiple DNA replication processes ^22, 23^. PARP1 activity is required for efficient SSBs repair and single-strand DNA gap-filling pathway in OFs during DNA replication ^21^. The DNA replication machinery is precisely controlled by cyclin-dependent kinases (CDKs), which phosphorylate several replication factors to allow them to enter the S phase and promote DNA synthesis ^24^. During replication fork progression, single-strand DNA is fragile and protected by replication protein A (RPA) ^25^. When replication forks stall, ataxia telangiectasia mutated and Rad3-related protein (ATR) is activated and phosphorylates CHK1 and RPA to resolve or eliminate the impediment, promoting replication fork recovery ^26, 27^. Furthermore, DNA damage, such as base damage, SSBs, DSBs, occurs upon genotoxic stress and replication errors, which lead to DNA replication stress, genomic instability, genetic mutation, and tumorigenesis.

In this study, we found that PNKP was required for the single-strand DNA gap-filling pathway during DNA replication. Defects in PNKP induce the accumulation of single-strand gaps in OFs and genome instability. We also found that CDKs phosphorylate PNKP on threonine 118 (T118), mainly in the S phase, and that CDK-mediated PNKP phosphorylation allows it to be recruited to single-strand DNA gaps on OFs via interaction with replication protein A 2 (RPA2). Moreover, PNKP enzymatic activity, especially phosphatase activity, is required for processing of the ends of single-strand gaps on OFs. Taken together, our data suggest that phosphorylation-mediated PNKP recruitment to single-strand DNA gaps on OFs and the end-processing activity of PNKP are critically important for preventing genome instability through appropriate regulation of DNA replication.

## Results

### Generation of PNKP knock out U2OS cell line

In a previous study, we found that the depletion of PNKP resulted in impaired cell growth in mice^28^. To confirm that this phenotype is observed in human cells, we initially generated PNKP knockout U2OS cells (*PNKP^−/−^*cells) using CRIPSR/Cas9 Nickase (D10A) targeting exon 4 of *PNKP* coding region ^29^. We obtained two clones (C1 and C2) that showed complete loss of PNKP, as confirmed by western blot analysis with both N-terminus and C-terminus PNKP-recognized antibodies and DNA sequencing (Fig. 1A and S1A and B). To confirm that these *PNKP^−/−^* cells have functional deficiencies in DNA repair, we analyzed repair abilities of DSBs and SSBs. *PNKP^−/−^*cells showed a delay in diminishing the phosphorylation of histone H2AX and KAP1, DSBs markers, after ionizing-radiation (IR) induced DNA damage, suggesting *PNKP^−/−^* cells defect DSB repair ability (Fig. S2A). A combination of a poly ADP-ribose glycohydrolase inhibitor (PARGi) and SSBs induction, such as via IR, can detect SSBs stained with anti-pan-ADP-ribose-binding reagents by immunofluorescence ^30^. We assessed SSB repair activity in *PNKP^−/−^* cells (Fig. S2B, C, D, and E). Meanwhile, non-treated *PNKP^−/−^* cells showed increasing ADP-ribose intensity, and IR-exposed *PNKP^−/−^*cells showed a marked loss of ADP-ribose intensity, indicating that *PNKP^−/−^*cells impaired the SSBs repair activity. Furthermore, *PNKP^−/−^*cells showed increased sensitivity to hydrogen peroxide (H_2_O_2_), a powerful inducer of SSBs, as well as IR (Fig. S2F and G). These findings validate the deficiency in SSBs and DSBs repair in *PNKP^−/−^*cells, aligning with the observations made in a previous study ^31^. Further, *PNKP^−/−^* cells showed increased sensitivity to the inhibition of DNA replication by hydroxyurea (HU) and camptothecin (CPT) treatment (Fig. S2H and I). Taken together, these results indicate that *PNKP^−/−^* cells were successfully established.

**Fig. 1.**
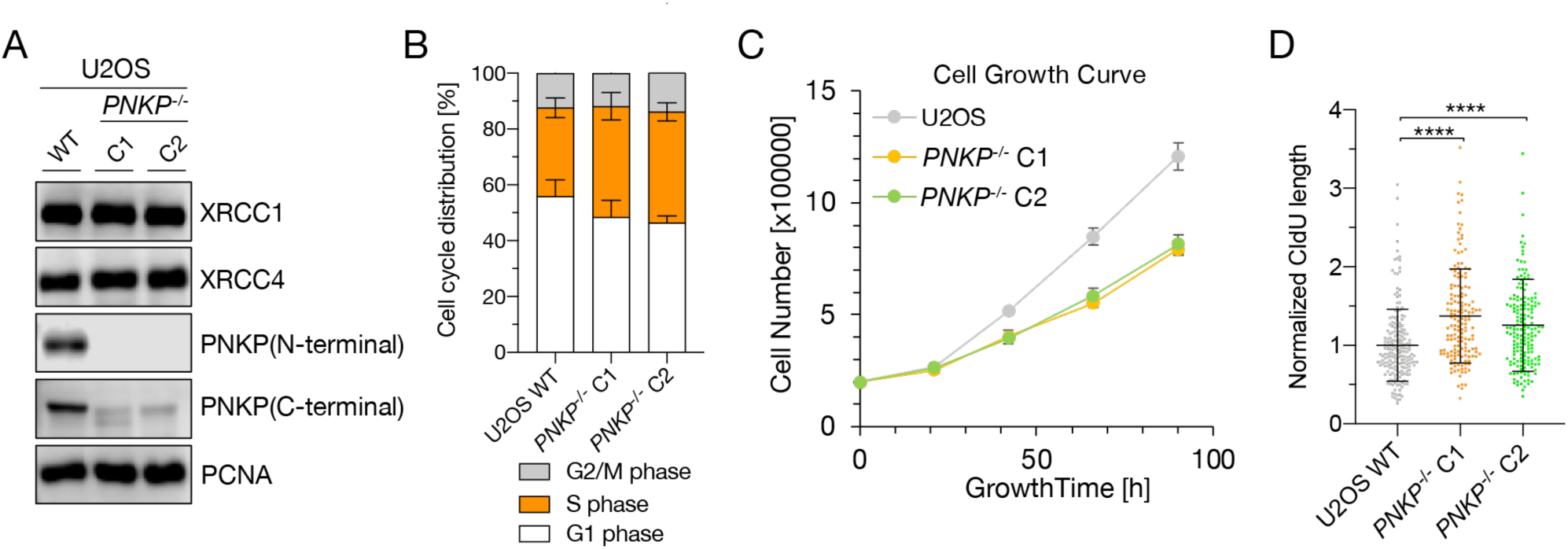
PNKP is important for proper S phase progression and cell proliferation. A: Protein expression analysis of PNKP knockout U2OS cells (*PNKP^−/−^*cells) generated by CRISPR/Cas9 genome editing. Protein expression of PNKP in *PNKP^−/−^* clone 1 (C1) and clone 2 (C2) were confirmed by western blotting with N-terminal and C-terminal recognized PNKP antibodies, respectively. XRCC1, XRCC4, and PCNA antibodies were used as loading control. B: Flowcytometric analysis for cell cycle distribution of U2OS WT and PNKP^−/−^ C1 and C2 cells. The cells were stained with Propidium Iodide (PI), and synthesized DNA was labeled by EdU. Percentage of each cell cycle is shown in vertical axis, cell types are shown in horizontal axis. C: Measurement of growth rate of U2OS WT and of *PNKP^−/−^* C1 and C2 cells. Cell numbers (shown in vertical axis) were counted at indicated time points (shown in horizontal axis). D: Quantified results of DNA fiber length in U2OS WT and *PNKP^−/−^* C1 and C2 cells. Nascent synthesized DNA was labeled by CldU. CIdU tract lengths in the indicated cell lines were plotted as scatter plots of fork speed combined from 3 independent experiments. >100 fibers per sample per experiment. Error bars represent standard deviation (SD). All experiments were performed at least three times independently. Statistical significance was assessed by one-way ANOVA with post hoc Sidak’s multiple comparisons test (\*\*\*\**P*<0.0001).

### PNKP is important for proper S phase progression and cell proliferation

Since cell proliferation is associated with DNA replication, we analyzed cell cycle distribution of *PNKP^−/−^* cells using flowcytometry, and the analysis revealed that *PNKP^−/−^* cells are accumulated in the S phase (Fig 1B and S3). *PNKP^−/−^* cells also exhibited slower cell growth than WT cells, which is consistent with our previous study in mice (Fig. 1C) ^28^. Next, we investigated whether PNKP is involved in replication fork progression using DNA fiber assays. Despite the slower cell growth, *PNKP^−/−^* cells had faster progression of DNA replication fork than WT cells (Fig. 1D). This result suggests that PNKP might be involved in the progression of DNA replication forks because a similar phenotype has been reported in PCNA polyubiquitination mutant (K164R) cells, which were unable to reduce the replication fork speed ^32^. PCNA KR mutant cells also showed slow cell proliferation because the accelerated replication speed failed to protect nascent DNA degradation. This resulted in a slower cell proliferation phenotype, reminiscent of that observed in *PNKP^−/−^* cells.

### PNKP phosphorylation, especially of T118, is important for proper S phase progression and cell proliferation

Subsequently, in order to identify the region of PNKP involved in fork progression, we measured the growth rate and the speed of fork progression in *PNKP^−/−^* cells transiently expressing PNKP deletion mutants (D1: FHA domain, D2: linker region, D3: phosphatase domain, and D4: kinase domain) (Fig. 2A-C, S4A) ^8^. D2 mutant-expressing cells showed slower proliferation than cells expressing WT PNKP and other mutants. Furthermore, D2 mutant-expressing cells also showed an increased speed of the replication fork compared to WT and D1 mutant-expressing cells, although D3 and D4 showed mildly high-speed fork progression. Since the linker region of PNKP is considered to be involved in fork progression, we attempted to identify essential amino acids for DNA replication in the linker region and identified five potential phosphorylation sites: serine 114, threonine 118, threonine 122, serine 126, and serine 143, using PhosphoSitePlus (Fig. 2D) ^33^.

**Fig. 2.**
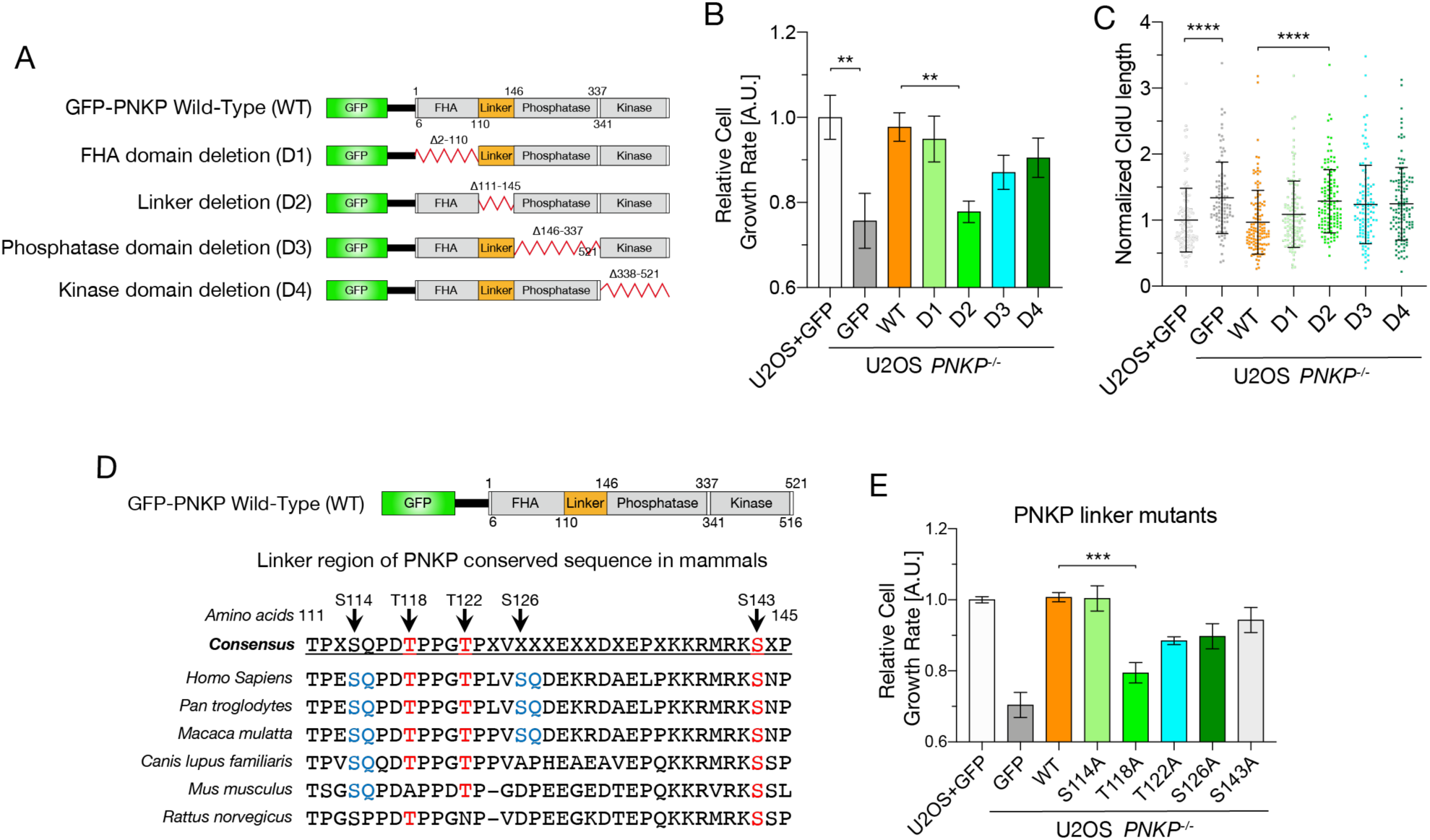
Phosphorylation of PNKP, especially on T118, is required for cell proliferation and DNA replication. A: Schematic diagrams of the structure of GFP -tagged human PNKP WT and deletion mutants (D1-D4). B: Cell growth rate in U2OS WT and PNKP deletion mutant expressing *PNKP^−/−^*C1 cells. Cell growth rate was normalized by GFP expressing U2OS cells. Error bars represent standard error of the mean (SEM). C: Quantified results of DNA fiber length in U2OS WT and *PNKP^−/−^* C1 cells expressing indicated PNKP deletion mutants. Nascent synthesized DNA was labeled by CldU. Error bars represent SD. D: Alignment of amino acid sequences in linker region among mammalian species. SQ/TQ motif (blue) is PI3 kinase substrate motif. TP (red) is CDK substrate motif. E: Cell growth rate in U2OS WT and *PNKP^−/−^* C1 cells expressing indicated point mutants. Cell growth rate was normalized by GFP expressing U2OS cells. Error bars represent SEM. All experiments were performed at least three times independently. Statistical significance was assessed by one-way ANOVA with post hoc Sidak’s multiple comparisons test (\*\**P*<0.01, \*\*\**P*<0.001, \*\*\*\**P*<0.0001).

Amino acids S114, T118, T122, and S143 are highly conserved among mammalian species. S114 and S126 form a typical SQ/TQ motif, and it has been reported that S114 is phosphorylated by ATM and S126 is phosphorylated by ATM and DNA-PKcs ^9, 34^. T118, T122, and S143 are novel phosphorylation sites, and T118 is predicted to be a CDK phosphorylation substrate motif (S/TP). To elucidate the importance of these amino acids in DNA replication, we constructed phosphorylation-mutant vectors containing five predicted phosphorylated amino acids substituted with alanine and measured the growth rates of these transfectants (Fig. 2E and S4B and C). T118A mutant-expressing cells exhibited a marked delay in cell growth, which was not observed for S114A, although T122A, S126A, and S143A were slightly delayed. These results suggest that the linker region of PNKP, especially the phosphorylation of PNKP T118, is required for S phase progression and proper cell proliferation.

### CDKs phosphorylate T118 of PNKP and pT118-PNKP interacts with nascent DNA on replication forks

To assess the importance of T118 phosphorylation in DNA replication, we generated antibodies that recognized the phosphorylated S114 (pS114), a control to confirm pT118 specificity in DNA replication, and phosphorylated T118 (pT118) peptides. Sufficient titers and specificities of the pS114 and pT118 antibodies were confirmed using ELISA (Fig. S5A, B, C, and D). The pT118 PNKP antibody was used for western blotting of lysates from U2OS WT-, green fluorescent protein (GFP)-PNKP WT-, or T118A-expressing cells. Unfortunately, this antibody cross-reacted with proteins of approximately 55 kDa, which is close to the apparent molecular mass of endogenous PNKP (Fig. S5E). Nevertheless, this antibody clearly recognized GFP-tagged PNKP (GFP-PNKP) but not GFP-PNKP T118A. Therefore, GFP-PNKP expression was used to examine T118 phosphorylation. To determine whether phosphorylation of T118 is DNA replication-specific, we synchronized HCT116 cells transiently expressing GFP-PNKP using a double thymidine block and released them at specific times (Fig. S6A). After synchronization at the indicated cell cycle phase, cells were extracted and used for western blotting (Fig. 3A). Because Cyclin A2 peaks at mid to late S/G2 phase and Cyclin E1 peaks at early S phase, we used these proteins as cell cycle markers ^35, 36,37^. pT118-PNKP was detected in asynchronized cells but increased particularly in the S phase, similar to Cyclin A2 expression levels. However, this effect was very weak during mitosis, suggesting that T118 phosphorylation plays a specific role in the S phase. Since amino acids around T118 contain CDK substrate motif, we treated cells with the CDK inhibitor roscovitine (CDKi) ^38, 39^ for 1 h. GFP-PNKP-expressing *PNKP^−/−^*cells showed decreased phosphorylation of T118 (Fig. 3B). In contrast, pS114 was not affected by CDKi treatment, indicating that CDKs specifically phosphorylate PNKP at T118 but not at S114. To identify the type of CDKs that phosphorylate PNKP at T118, we measured the direct phosphorylation activity using purified CDKs, cyclin, and PNKP, and detected them using the T118P PNKP antibody (Fig. 3C). CDK1/Cyclin A2 and CDK2/Cyclin A2 markedly phosphorylated PNKP, whereas CDK4/Cyclin D1 and CDK2/Cyclin E1 phosphorylated PNKP to a lesser extent, suggesting that CDK1/CyclinA2 and CDK2/CyclinA2 complexes are potential kinases of PNKP T118. We investigated the phosphorylation levels of PNKP at T118 under co-overexpression of Cyclin A2 and CDK1 and 2 (Fig. 3D). Overexpression of both CDK2/Cyclin A2 and CDK1/Cyclin A2 showed an increased phosphorylation level at T118, suggesting that CDK1/cyclinA2 and CDK2/CyclinA2 complexes are potential kinases of PNKP T118.

**Fig. 3.**
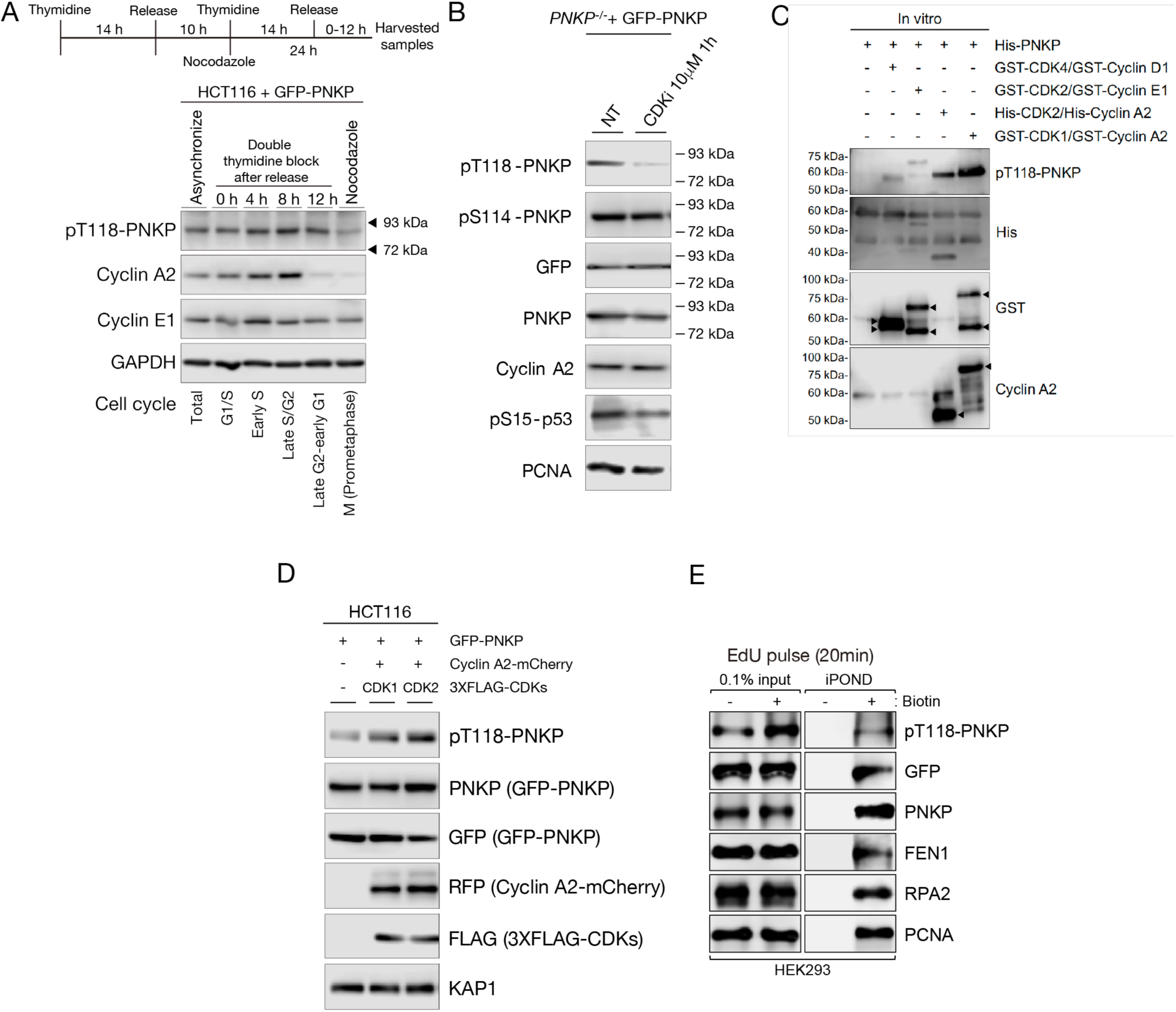
CDKs phosphorylate T118 of PNKP and pT118-PNKP interacts with nascent DNA on replication forks. A: Scheme and protein expression levels of GFP-PNKP expressing HCT116 cells after release from double thymidine block. After released, cells were collected at indicated time points, and used for western blotting. pT118 PNKP antibody was generated for this study. Cyclin A2 and E1 antibodies were used for cell cycle markers. GAPDH antibody was used as a loading control. B: T118-PNKP phosphorylation levels of GFP-PNKP expressing *PNKP−/−* C1 cells after treatment of 10 μM CDKs inhibitor, roscovitine, for 1h. Cells were lysed and applied for western blotting. GFP and PNKP antibodies were used for confirming GFP-PNKP expression levels. Cyclin A2 antibody was used as a cell cycle marker. pS15-p53 antibody was used for confirming CDKs inhibition. pS114-PNKP antibody, generated in this study, was used for confirming pT118 specificity against CDKs inhibition. PCNA antibody was used as a loading control. C: In vitro analysis of PNKP phosphorylation on T118 by CDK/Cyclin complex. Purified His-PNKP and each CDK/Cyclin complex were incubated with reaction mixture and detected with western blotting by T118P PNKP, His, GST, and Cyclin A2 antibodies. D: HCT116 cells were lysed at 5 days and 3 days after transfection with GFP-PNKP and co-transfection with mCherry2-Cyclin A2, and 3XFLAG-CDK1 or CDK2, respectively, and applied for western blotting. pT118 PNKP level was analyzed by pT118 PNKP specific antibody. PNKP expression was analyzed with PNKP or GFP antibodies. Cyclin A2 expression was analyzed with RFP antibody. KAP1 antibody was used as loading control. E: Analysis of isolated proteins from nascent DNA using iPOND technique. Proteins bound to EdU labeled DNA in GFP-PNKP expressing HEK293 cells were isolated using click reaction with biotin-azide followed by streptavidin-pulldowns and detected by western blotting. 0.1% of lysate used in Streptavidin-pulldowns represented as 0.1% input. All experiments were performed at least three times independently.

To elucidate the role of PNKP T118 phosphorylation in DNA replication, we isolated proteins from nascent DNA (iPOND) to confirm the interactions between PNKP and nascent DNA on replication forks (Fig. S6B). After 5-ethynyl-2’-deoxyuridine (EdU) incorporation for 20 min in GFP-PNKP-expressing HEK293 cells, proteins bound to nascent DNA were extracted and detected by western blotting (Fig. 3E). We found that not only the replication fork-associated proteins, FEN1, RPA2, and PCNA, but also WT-PNKP and T118-phosphorylated PNKP interacted with nascent DNA. Taken together, these results suggest that the CDK1/Cyclin A2 or CDK2/Cyclin A2 complex potentially regulates the phosphorylation level of PNKP T118 in the S phase to interact with nascent DNA.

### Phosphorylation of PNKP at T118 is required for preventing single-strand DNA gap formation between Okazaki fragments

Considering that one of the known targets of PNKP in DNA repair is SSBs, including gapped DNA structures, we anticipated that PNKP might be involved in the formation of single-strand DNA nicks and/or small gaps during DNA replication, not only in the repair of exogenous DNA damage. To confirm this hypothesis, we used HU and Emetine (EME), a DNA replication inhibitor that blocks single-strand gap formation on replication forks via proteosynthesis inhibition ^40, 41^. We analyzed the extent of single-strand DNA nicks and/or gaps by poly ADP-ribosylation in *PNKP^−/−^* cells (Fig. 4A). HU treatment slightly increased the amount of poly ADP-ribosylation, whereas EME prevented poly ADP-ribosylation by inhibiting single-strand nick and/or gap formation in the S phase of U2OS WT cells. These results were consistent with those of a previous study ^21^. Although *PNKP^−/−^* cells showed a high level of ADP-ribose intensity spontaneously, HU and EME treatment reduced the ADP-ribose intensity, albeit to a slightly higher extent than WT cells, suggesting that loss of PNKP leads to an accumulation of single-strand DNA nicks and/or gaps structures via DNA replication. To specifically investigate the extent of single-strand DNA nicks and/or gaps in the S phase, we transiently expressed WT PNKP or T118A in *PNKP^−/−^*cells and incubated them with EdU-containing medium to separately assess ADP-ribose intensity inside (EdU-positive) or outside (EdU-negative) the S phase (Fig. 4B and C). In the S phase, T118A expressing cells showed high ADP ribose intensity, similar with *PNKP^−/−^* cells, and WT PNKP rescued this effect, suggesting that PNKP function and T118 phosphorylation are required for gap-less DNA replication. To investigate the origin of single-strand DNA nicks and/or gaps, we used a flap endonuclease 1 inhibitor (FEN1i), which prevents the resection of overhanging nucleotides from the ends of OFs, leading to unligated OFs ^42, 43, 44^. We assessed the ADP-ribose intensity of WT PNKP or T118A-expressing *PNKP^−/−^*cells treated with either DMSO or FEN1i only during the S phase (EdU-positive cells) (Fig. 4D and E). *PNKP^−/−^* cells and T118A-expressing cells strongly increased the intensity of ADP ribose compared with U2OS- and PNKP WT-expressing cells in the S phase even after DMSO treatment. FEN1i-treated U2OS and PNKP WT-expressing cells showed higher ADP-ribose intensity than DMSO-treated cells. In contrast, FEN1i treated *PNKP^−/−^*or T118A-expressing cells eliminated the disparity in ADP-ribose intensity, suggesting that PNKP suppresses the OFs-mediated gap formation in an epistatic pathway with FEN1. We subsequently assessed the implications of OFs maturation in replication fork progression using a DNA fiber assay (Fig. 4F). FEN1i treatment induced high-speed fork progression, although HU treatment led to stalled forks, suggesting that inhibition of OFs maturation by FEN1i leads to defects in replication fork protection. We analyzed this phenotype in cells expressing *PNKP^−/−^*, T118A, and T118D (a phosphor-mimetic mutant) (Fig. 4G). *PNKP^−/−^* and T118A-expressing cells showed spontaneous fast fork progression, and FEN1i treatment did not further accelerate the DNA replication speed, indicating that the T118A mutation is enough to provoke faster fork progression due to unligated OFs generation, as seen with FEN1 inhibition. Moreover, T118D-expressing cells showed normal DNA replication speed, similar to that of PNKP WT-expressing cells, and FEN1i treatment accelerated the speed of DNA replication in T118D-expressing cells, suggesting that PNKP and its phosphorylation at T118 play essential roles in OFs maturation and accurate DNA replication. Additionally, to address whether phosphorylation of T118 promotes its recruitment to gapped DNA structures, we assessed a binding ability to a single-strand DNA gap structure using WT PNKP, T118A, and T118D mutants expressing *PNKP^−/−^* cell lysates (Fig. 4H). The T118A mutant showed impaired gapped DNA-binding ability, whereas the WT PNKP and T118D mutant showed relatively higher binding ability than the PNKP T118A mutant. Taken together, we conclude that the phosphorylation of PNKP at T118 is required for its recruitment to single-strand DNA nicks and/or gaps to prevent the eruption of unligated OF-mediated post-replicative single-strand DNA gap formation during DNA replication.

**Fig. 4.**
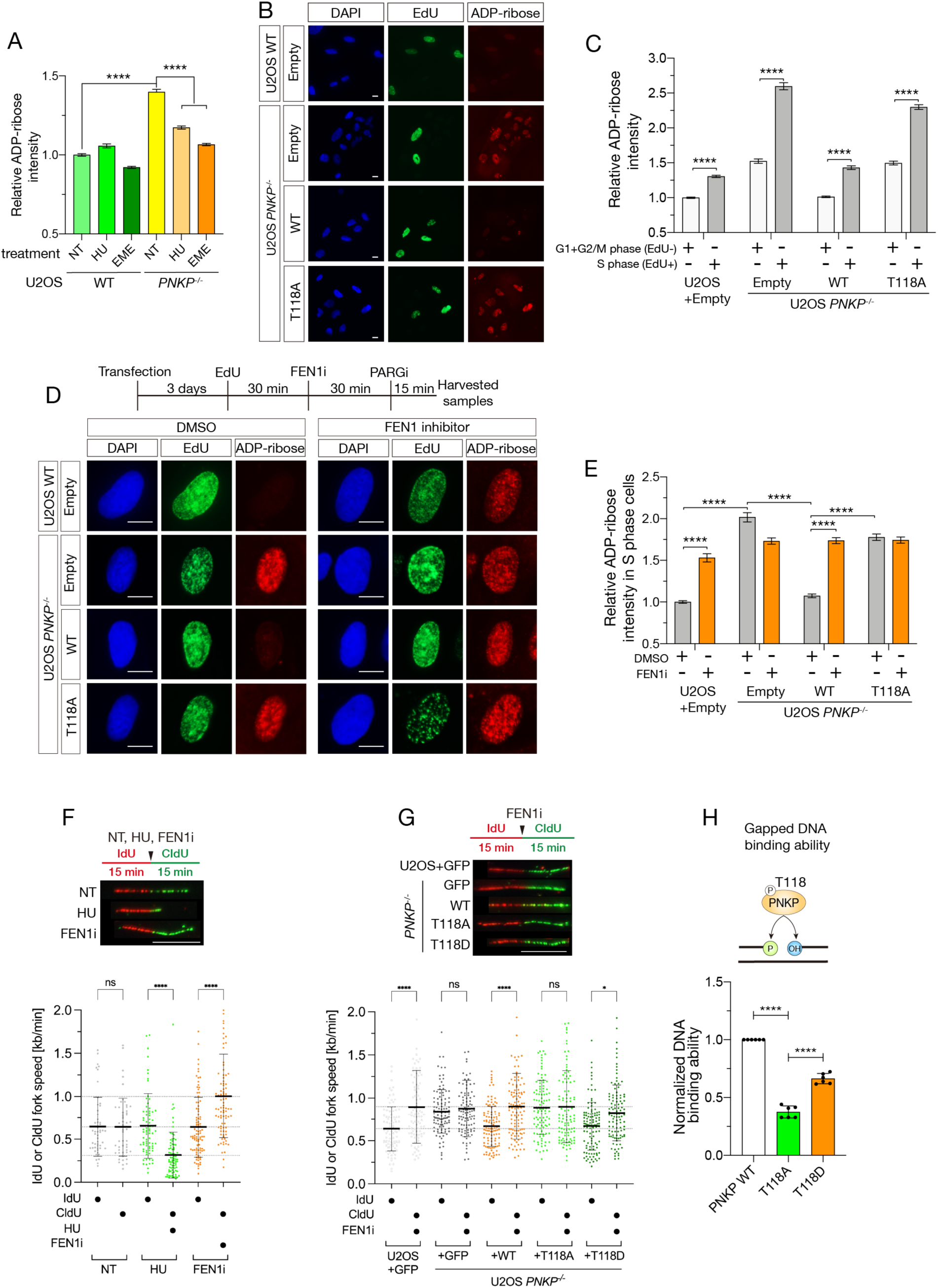
Phosphorylation of PNKP on T118 is required for preventing single-strand DNA gap formation between Okazaki fragments. A: Measurement of SSBs, detected by ADP-ribose intensity, in U2OS WT and *PNKP^−/−^* C1 cells under Hydroxyurea (HU) or Emetine (EME) treatment. Cells were pre-treated with poly ADP-ribose glycohydrolase inhibitor (PARGi) for 30 min prior to the treatment of HU or EME. ADP-ribose intensities were normalized by the intensity of non treated (NT) U2OS. Error bars represent SEM. B and C: Representative images (B) and quantified results (C) of measurement of ADP-ribose intensity in U2OS WT and PNKP WT, PNKP T118 expressing *PNKP^−/−^* C1 cells. Synthesized DNA was labelled by EdU and EdU-positive cells were defined as S phase (grey) and the other cells were defined as G1+G2/M phase (white). Error bars represent SEM. D and E: Scheme, representative images (D) and quantified results (E) of the experiments for measurement of ADP-ribose intensity in U2OS WT and *PNKP^−/−^* C1 cells expressing GFP, PNKP WT, and PNKP T118A mutant under dimethyl sulfoxide (DMSO) (negative control) and Flap endonuclease 1 inhibitor (FEN1i) treatment only in EdU-positive (S phase) cells. Error bars represent SEM. F: Measurement of DNA synthesis speed in U2OS WT cells under NT, 2 mM HU and 10 μM FEN1 inhibitor treatment analyzed by DNA fiber assay. At least 100 DNA fibers were measured. Error bars represent SD. G: Measurement of DNA synthesis speed in U2OS WT and PNKP^−/−^ C1 cells expressing GFP (con), PNKP, T118A, or T118D mutants under 10 μM FEN1 inhibitor treatment analyzed by DNA fiber assay. At least 100 DNA fibers were measured. Error bars represent SD. H: Measurement of the gapped DNA binding ability of PNKP WT, T118A, and T118D mutants. Nuclear extracts were harvested from U2OS *PNKP^−/−^* C1 cells at 2 days after transfection with GFP-PNKP WT, T118A or T118D mutants. GFP antibody was used to detect GFP-PNKPs bound to the gapped double-stranded DNA. Error bars represent SEM. In all panels, scale bar indicates 10 μm. All experiments were performed at least three times independently. Statistical significance was assessed by one-way ANOVA with post hoc Sidak’s multiple comparisons test (ns = no significant, \**P*<0.1, \*\*\*\**P*<0.0001).

### Phosphatase activity of PNKP is important for the end-processing of Okazaki fragments

The main function of PNKP is to catalyze 5’-phosphorylation and 3’-dephosphorylation of the DNA ends. To clarify the role of these PNKP enzymes, we constructed phosphatase-dead (D171A) and kinase-dead (K378A) PNKP mutants ^3, 14, 45^. We first attempted to establish stable clones expressing both mutants; however, we could not obtain stable phosphatase-dead (D171A) clones. This is consistent with previous observations, in which PNKP-mutated MCSZ patient cells, exhibiting mutations in the phosphatase domain, showed unstable expression of PNKP ^11^, and recombinant phosphatase-dead PNKP was unstable ^15^. Therefore, these mutants were expressed transiently in *PNKP^−/−^* cells, and protein expression was assessed through western blotting (Fig. 5A). First, we assessed the phosphatase and kinase enzymatic activities of the PNKP T118A mutant and performed biochemical assays using PNKP mutants expressing cell extracts (Fig. 5B and S7). Fluorescence-labeled single strand break gap oligo DNA was mixed with cell lysates extracted from U2OS WT or *PNKP^−/−^* cells expressing several PNKP mutants. The D171A and K378A mutants were used as phosphatase-dead and kinase-dead controls, respectively. Although the phosphatase-dead mutant showed slightly lower kinase activity and the kinase-dead mutant showed lower phosphatase activity than WT PNKP, it is possible that each mutant is structurally unstable and affects the activity of the enzyme. Intriguingly, T118A PNKP was still capable of dephosphorylating and phosphorylating gapped DNA ends *in vitro*, albeit to a lesser extent than WT PNKP. These results suggest that phosphorylation of PNKP at T118 is required for its recruitment to the gapped DNA structure but not directly for its enzymatic activity.

**Fig. 5.**
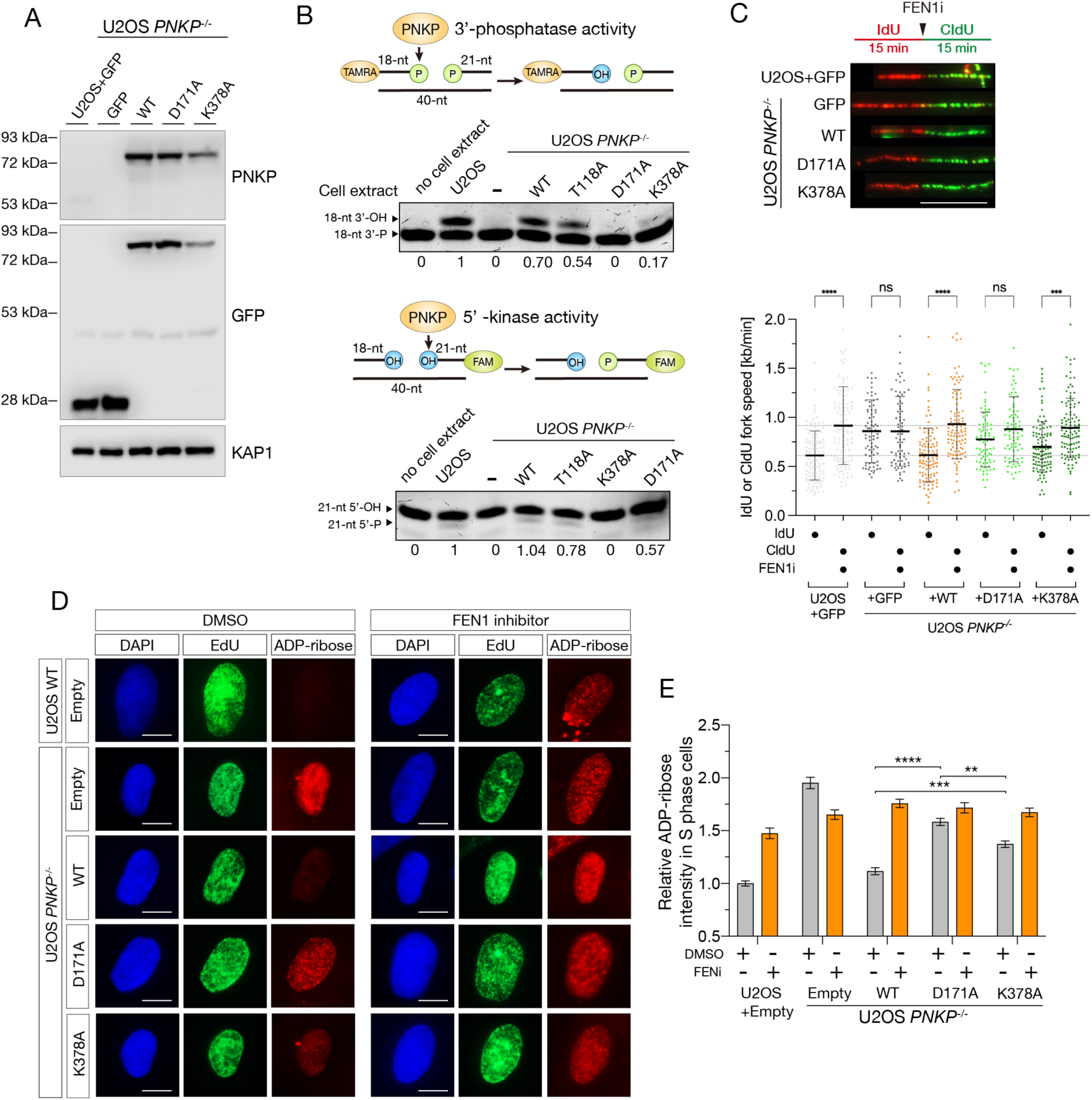
Phosphatase activity of PNKP is important for the end-processing of Okazaki fragments. A: Protein expression analysis for PNKP phosphatase-dead (D171A) and kinase-dead (K378A) mutants in U2OS WT and *PNKP^−/−^* C1 cells expressing GFP, PNKP WT, D171A and K378A mutants confirmed by western blotting. KAP1 antibody was used as a loading control. B: PNKP phosphatase and kinase activity biochemical analysis. Indicated cell extracts harvested from U2OS WT and *PNKP^−/−^* C1 cells expressing PNKP WT, T118A, D171A (phosphatase dead), and K378A (kinase dead) were incubated with TAMRA or 6-FAM labelled oligonucleotide duplex harboring a SSB GAP structure. Arrows indicate the positions of the TAMRA labelled phosphatase substrates (left) and 6-FAM labelled kinase substrates (right). Band intensity of 18-nt 3’ -OH (left) and 21-nt 5’ -P (right) were analyzed using Image J software and indicated. C: Measurement of the speed of DNA synthesis in U2OS WT and *PNKP^−/−^*C1 cells expressing GFP (con), PNKP, D171A and K378A mutants under 10 μM FEN1 inhibitor treatment analyzed by DNA fiber assay. At least 100 DNA fibers were measured. Error bars represent SD. D and E: Representative images (D) and quantified results (E) of the experiments for measurement of ADP-ribose intensity in U2OS WT and *PNKP^−/−^* C1 cells expressing GFP, PNKP WT, PNKP D171A and PNKP K378A mutants under DMSO (negative control) and FEN1i treatment only in EdU-positive (S phase) cells. Error bars represent SEM. In all panels, scale bar indicates 10 μm. All experiments were performed at least three times independently. Statistical significance was assessed by one-way ANOVA with post hoc Sidak’s multiple comparisons test (ns = no significant, \*\**P*<0.01, \*\*\**P*<0.001 \*\*\*\**P*<0.0001).

Next, we analyzed the rate of fork progression in cells expressing these mutants using a DNA fiber assay (Fig. 5C). Interestingly, D171A expressing cells showed a spontaneously higher speed of fork progression than WT PNKP-expressing cells, and FEN1i treatment did not accelerate DNA replication. However, K378A-expressing cells showed slightly more spontaneous faster fork progression than WT PNKP-expressing cells, suggesting that PNKP enzymatic activities, especially those of phosphatase, are required for accurate fork progression and protection. Subsequently, we elucidated whether the end-processing activities of PNKP are important for OF-mediated single-strand DNA gap formation. These PNKP mutants expressing *PNKP^−/−^* cells were treated with FEN1i. After which, we measured the ADP-ribose intensity in the S phase (EdU-positive cells) (Fig. 5D and E). D171A showed a high ADP-ribose intensity, while K378A showed a relatively high ADP-ribose intensity without FEN1i treatment. Moreover, FEN1i-treated cells showed high levels of ADP-ribose intensity in all cells. Taken together, these results suggest that PNKP phosphatase and kinase activities, especially those of phosphatase, play an important role in the end processing of OFs, resulting in the suppression of OFs-mediated post-replicative single-strand DNA gap formation and accurate DNA replication.

### PNKP is involved in postreplicative single-strand DNA gap-filling pathway

Upon deficient ligation of OFs, post-replicative single-strand DNA gaps evolve from unligated OFs, which are sensed by PARP1 and repaired via the PARP1-dependent gap-filling pathway for OFs maturation ^46^. The PARP1-dependent gap-filling pathway is associated with XRCC1, a binding scaffold protein of PNKP in the single-strand break repair pathway ^21^. To determine whether PNKP plays a critical role in the PARP-dependent gap-filling pathway, we performed a DNA fiber assay (Fig. 6A). Cells were labeled with 5-iodo-2’-deoxyuridine (IdU) for 15 min, followed by 60 min of 5-chloro-2’-deoxyuridine (CldU) labeling in the presence or absence of an FEN1 inhibitor and/or PARP inhibitor, and the individual tract lengths of nascent DNA labeled with IdU or CldU were quantified. Where indicated, labeled DNA was treated with an endonuclease, S1 nuclease, to detect post-replicative single-strand DNA gaps (Fig. 6B, C, and D). PNKP*^−/−^* cells showed spontaneously faster progression of replication measured by IdU or CldU labeled DNA tract lengths. Interestingly, in untreated *PNKP^−/−^*cells, nascent DNA was cleaved upon S1 nuclease treatment, similar to the response observed in U2OS WT cells subjected to both FEN1i and PARPi treatments. Notably, U2OS WT cells treated with FEN1i or PARPi individually did not exhibit this phenotype. These results suggest that PNKP is involved in both single-strand DNA gaps, such as FEN1-related OFs ligation, and the PARP1-dependent gap-filling pathway.

**Fig. 6.**
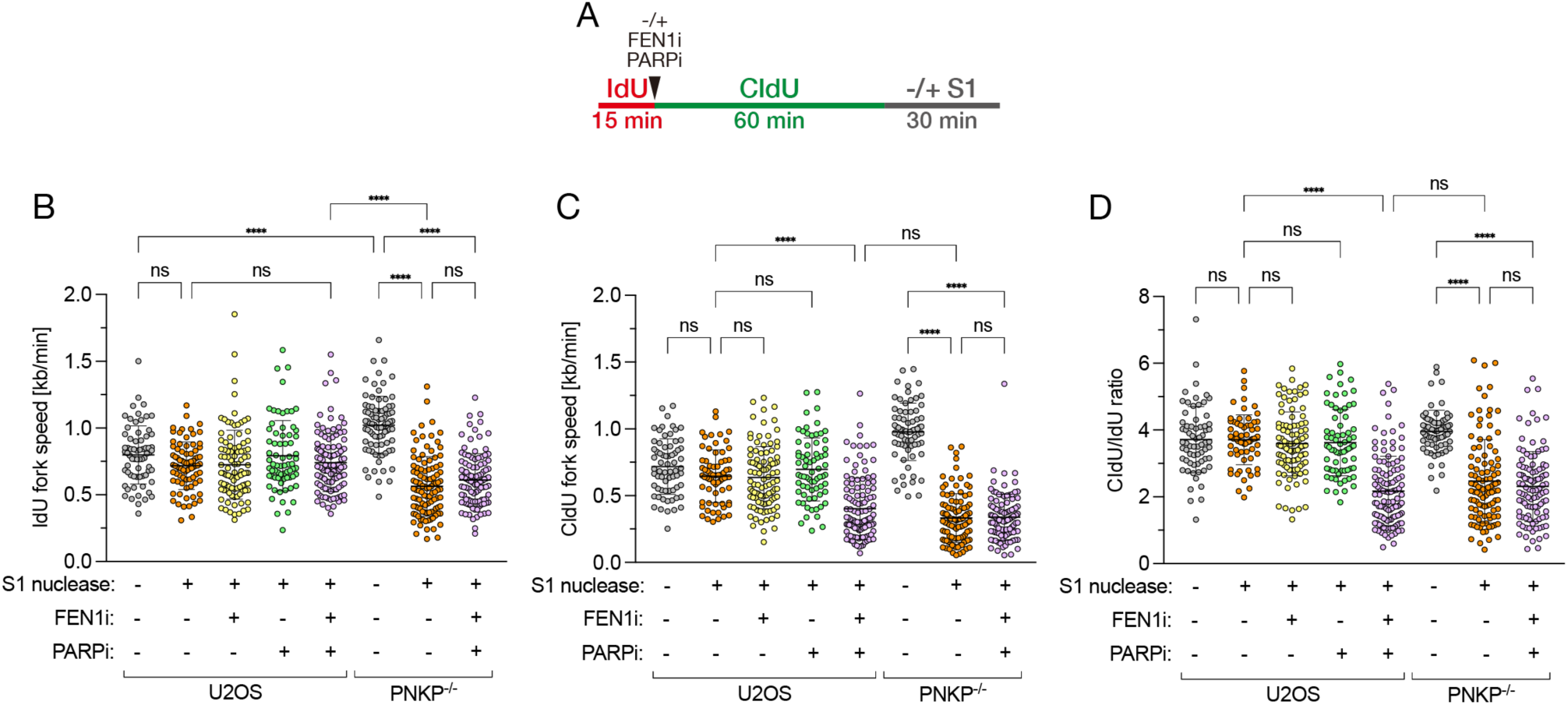
PNKP is required for repairing postreplicative single-strand DNA gaps derived from unligated Okazaki fragments. A: Schematic for measuring the replication fork progression in U2OS WT and *PNKP^−/−^* C1 cells with 10 μM FEN1 inhibitor and/or 10 μM PARP inhibitor treatment followed by S1 nuclease treatment. B, C, and D: CldU or IdU tract lengths in dual-labeled DNA fibers in the indicated cell lines and treatments were measured. The speed of CldU- or IdU-labeled replication fork progression was calculated assuming a constant stretching factor of 2 kb/μm ^46^.Error bars represent SD. All experiments were performed at least three times independently. Statistical significance was assessed by one-way ANOVA with post hoc Sidak’s multiple comparisons test (ns = no significant, \*\*\*\**P*<0.0001).

### Phosphorylation of PNKP at T118 is essential for genome stability

Finally, we investigated whether the function of PNKP in DNA replication via the phosphorylation of PNKP at T118 is required for genome stability. PNKP T118A-expressing *PNKP^−/−^*cells showed increasing frequency of micronuclei and chromosome bridges compared with the WT PNKP complemented cells without any treatment as well as with IR or HU treatment (Figure 7A and B). Furthermore, we assessed the SSBs and DSBs repair abilities of PNKP T118A-expressing cells based on the levels of poly-ADP ribosylation and γH2AX foci formation after IR (Fig. S8A, B and C). PNKP T118A-expressing cells showed high levels of ADP-ribose intensity and an increased number of γH2AX foci-positive cells, suggesting that T118 phosphorylation is essential for both SSB and DSB repair. Taken together, our results suggest that the loss of PNKP or T118 phosphorylation might cause endogenous single-strand DNA gap formation during DNA replication as well as repair endogenous DNA damage to prevent genome instability.

**Fig. 7.**
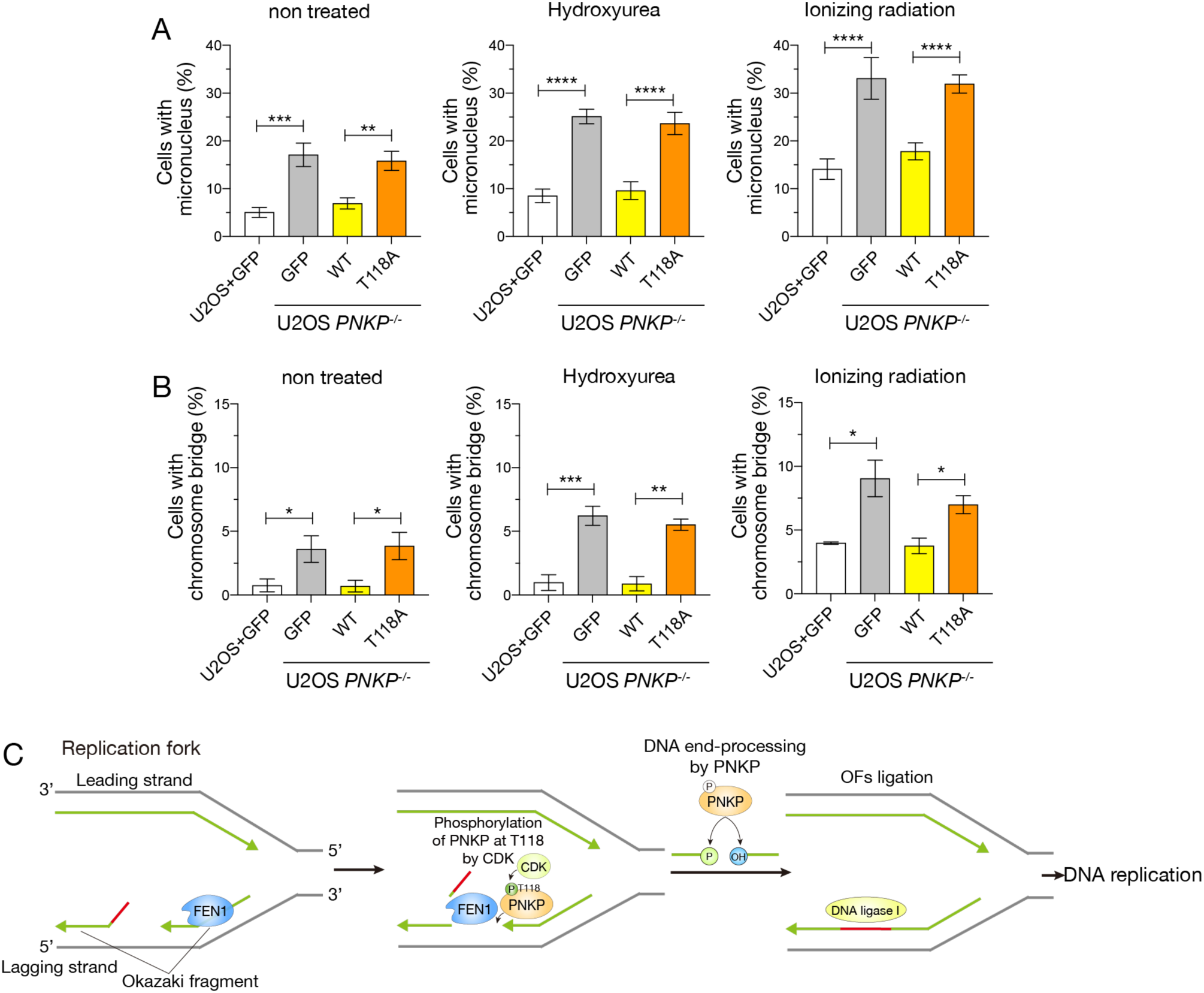
Phosphorylation of PNKP at T118 is required for genome stability. A and B: Measurement of the formations of micronuclei and chromosome bridges in U2OS WT and *PNKP^−/−^* C1 cells expressing GFP, PNKP WT, and PNKP T118A mutant exposed to 5 Gy IR or 2 mM HU. DNA were stained by DAPI at 24 h after each treatment. Cells with micronucleus (A) and chromosome bridge (B) were counted and graphed. At least 200 cells were counted. Error bars represent SEM. C: Schematic mechanisms of the involvement of PNKP in DNA replication, especially in end-processing of Okazaki fragments (OFs) and gap-filling pathway. T118 of PNKP is phosphorylated by CDKs for the recruitment to single-strand DNA nicks and/or gaps between OFs. OFs ends are processed by PNKP and become the ligatable ends prior to the OFs maturation. In PNKP-deficient, unligated OFs become postreplicative single-strand DNA gaps, and the gaps are sensed by PARP1 for proceeding PARP-dependent gap-filling pathway which required SSB repair proteins, XRCC1, LIG3 and PNKP. Impaired PARP-dependent gap filling leads to the high-speed DNA synthesis and the genome instability. All experiments were performed at least three times independently. Statistical significance was assessed by one-way ANOVA with post hoc Sidak’s multiple comparisons test (ns = no significant, \**P*<0.1, \*\**P*<0.01, \*\*\**P*<0.001, \*\*\*\**P*<0.0001).

## Discussion

In this study, we identified the phosphorylation of PNKP at T118 mediated by CDKs, potentially by the CDK1/CyclinA2 or CDK2/CyclinA2 complex, which is important for recruitment to the gapped DNA structure, including nicks between OFs and OF-mediated post-replicative single-strand DNA gap structures (Fig. 7C). Defects in PNKP phosphorylation at T118 result in the accumulation of single-strand DNA nicks and/or gaps during DNA replication due to a deficiency in OFs ligation and the subsequent gap-filling pathway.

PNKP consists of four regions: the FHA, linker, phosphatase, and kinase domains. Compared to other domains, the role of the linker region is still poorly understood, although it includes several residues that may be post-translationally modified. DNA damage signaling and DNA replication progression are often regulated by protein modifications such as phosphorylation and ubiquitination ^47, 48, 49^. Lysine is the main target of E3 ubiquitin ligase ^50^, and there is a clustered lysine region (137–142) in the PNKP linker region. However, we reported that this region includes a nuclear localization signal (NLS) and that alanine substitution prevented it from being transported to the nucleus ^7^. Therefore, we focused on regions other than amino acids 137–142. In the present study, we identified five predicted phosphorylation sites (S114, T118, T122, S126, and S143) in the linker region. Using laser microirradiation analysis, we assessed the accumulation of these PNKP mutants at DNA damage sites (Fig. S9). Although the T122A, S126A, and S143A mutants showed no significant differences from WT PNKP (Fig. S9A, B, and C), the mutants showed slightly reduced cell proliferation (Fig. 2E), suggesting that these residues may be involved in other cellular functions. In contrast, the S114A and T118A mutants showed a considerably attenuated accumulation of DNA damage sites. In addition, the phosphorylation mimic mutants, S114D and T118D, accumulated at DNA damage sites (Fig. S9D, E, F, and G). S114A mutant-expressing cells showed the presence of SSBs repair and normal cell growth (Fig. 2E and S7), suggesting that the phosphorylation of PNKP at S114 is likely to be important for DSBs repair. In contrast, the T118A mutants showed the absence of both SSBs and DSBs repair (Fig. S7). However, residual SSBs often cause DNA replication-coupled DSBs formation, and the impaired DSB repair ability of the T118A mutants may be due to a combination of impaired SSB repair ability and DNA replication-coupled DSBs formation.

Regulating the speed of DNA synthesis is important for accurate DNA replication and fork integrity ^51, 52^. *PNKP^−/−^* and T118A-expressing cells showed high-speed of DNA synthesis, resulting in slower cell proliferation and genome instability (Fig. 2, 4, 5, and 6). These observations are consistent with those in PCNA KR mutant cells and PARPi-treated cells ^32, 53^. We also found that enzymatic activity, especially the phosphatase activity of PNKP, is required for the end-processing of OFs during DNA replication (Fig. 5). *PNKP^−/−^* cells and mutant-expressing cells strongly increased the ADP-ribose intensity in the S phase even without FEN1i treatment, although FEN1i treatment in these cells leads to attenuated or diminished increase in the ADP-ribose intensity compared with WT PNKP-expressing cells, suggesting that PNKP acts as an epistatic pathway in FEN1 for OFs ligation (Fig. 5D and E). FEN1 catalyzes the removal of RNA/DNA fragments from OFs on the lagging strand. Our data also suggested that both the end-processing activity of PNKP and the exonuclease activity of FEN1 were required for OFs ligation (Fig. 4D and E, 5D and E, and 6). This observation is consistent with that of HU- and EME-treated *PNKP ^−/−^* cells showing decrease in the ADP-ribose intensity (Fig. 4A). Since EME inhibits single-strand gap formation ^40, 41^ and HU treatment increases the amount of OF-like DNA fragments ^54, 55, 56^, these reagents inhibit mature OF structures as PNKP-appropriate substrates. Moreover, we found that the phosphorylation of PNKP at T118 was regulated in a cell cycle-dependent and CDK-associated manner (Fig. 3A-C). Because both CDK1/Cyclin A2 and CDK2/Cyclin A2 are involved in PNKP phosphorylation, Cyclin A2 is likely important for these activities. We concluded that phosphorylation of PNKP at T118 allows it to be recruited to the ends of OFs, including single-strand nicks and/or gaps on the lagging strand in the S phase, and processes the ends for subsequent ligation. Furthermore, PNKP is required for the PARP1-dependent single-strand gap-filling pathway when unligated OFs are transformed into post-replicative single-strand DNA gap structures (Fig. 6).

In summary, because the maintenance of genome stability is central to life, the function of PNKP in DNA repair and OFs maturation is important for biological development. Although PNKP mutations cause several inherited diseases (MCSZ, AOA4, and CMT2B2) with neural developmental failure and neurodegeneration, almost all mutations are found in the FHA, phosphatase, or kinase domains and not in the linker region. This may be explained by the fact that mutations in the phosphorylated residue in the linker region are embryonic lethal. Since the inhibition of PNKP activity is an important target for anticancer drugs, in addition to the inhibition of phosphatase and kinase enzymatic activity ^57^, the inhibition of phosphorylation on T118 of PNKP might be a potent target for cancer therapy. Ultimately, this study presents a novel role for PNKP in processing the ends of OFs and the PARP-dependent single-strand gap-filling pathway during DNA replication, which may contribute to the elucidation of the biological basis of DNA replication and increase our understanding of the mechanisms underlying the onset of inherited diseases.

## Materials and Methods

### Cell culture

The human embryonic kidney cell line HEK293, the human colon cancer cell line HCT116, and the human osteosarcoma cell line U2OS were obtained from the American Type Culture Collection (ATCC), and U2OS *PNKP^−/−^* cell lines were established in this study. All cell lines were maintained in Dulbecco’s modified Eagle’s medium (DMEM; Nacalai Tesque Inc.) supplemented with 10% v/v fetal bovine serum (FBS; Hyclone, GE Healthcare) and penicillin/streptomycin (Nacalai Tesque Inc.) at 37 °C in humidified atmosphere containing 5% CO_2_ conditions. All cell lines were tested for *Mycoplasma* contamination using the e-Myco^TM^ Mycoplasma Detection PCR Kit (iNtRON Biotechnology, Inc., cat# 25235).

### Construction of plasmid DNA and mutagenesis

The pEGFP-C1 plasmid was purchased from Clontech. Full-length human PNKP cDNA was obtained by PCR from the cDNA pool of U2OS cells and inserted into the pEGFP-C1. Mutations were introduced using the PrimeSTAR mutagenesis basal kit (Takara Bio, cat# R046A) according to the manufacturer’s instructions. All the DNA constructs were verified by DNA sequence analysis. All primers for mutagenesis were designed using the Agilent QuikChange primer design program and are listed in Supplementary Table 1.

### cDNA and siRNA transfection

For cDNA transfection, PEI-MAX (Polysciences, Inc., cat# 24765) or Lipofectamine 3000 (Invitrogen, Thermo Fisher Scientific) were used according to the manufacturer’s instructions. For siRNA transfection (treatment time is typically–48–72 h), Lipofectamine RNAiMAX (Invitrogen, Thermo Fisher Scientific) was used according to the manufacturer’s instructions. All the siRNAs were used at a final concentration of 50 nM. The targeting sites and sequences of the siRNA oligonucleotides are listed in Supplementary Table 2.

### Genome editing by CRISPR/Cas9 system and establishment of PNKP^−/−^ cell lines

The pSpCas9n(BB)-2A-Puro (PX462) vector was purchased from Addgene. For the establishment of *PNKP*^-/-^ cells using CRISPR/Cas9 system, the sgRNA target sequences were cloned into the pSpCas9n(BB)-2A-Puro (PX462) vector and verified by DNA sequencing. U2OS cells were transfected with the targeting vectors and incubated for 2 days before the addition of selective medium containing 1.5 mg/mL puromycin (InvivoGen, cat# ant-pr-1). After 5 days, clonal cells were isolated by limiting dilution in 96-well plates. PNKP expression in single clones was analyzed by western blotting. Genomic mutations in both PNKP alleles present in U2OS cells were verified by DNA sequencing of a PCR-amplified genomic fragment cloned into a pEGFP-C1 vector. The sgRNA target sequences are shown in Supplementary Fig. 1.

### Laser microirradiation and live-cell imaging

U2OS cells were grown on glass-bottom 35-mm dishes (Matsunami Glass Ind., Ltd.), and the indicated GFP-tagged cDNA was transfected into the cells at 2 days before observation using the method described above. On the day after transfection, the culture media containing the transfection reagents were replaced with normal culture media without transfection reagents. On the day of observation, culture media were replaced with phenol red-free DMEM (Nacalai Tesque Inc.) supplemented with 10% v/v FBS, and 0.5 mg/mL Hoechst 33258 (Sigma-Aldrich) was added for the enhancement of the DNA damage at 30 min prior to the observation. A Leica TCS SP8 LIGHTNING Confocal Microscope (Leica Microsystems) with a 63 x /1.40 oil immersion objective lens was used for inducing localized DNA damage through laser microirradiation, live-cell observation, and image capture ^58, 59^. For quantification, 20 cells were irradiated with the laser, and the green fluorescence intensity was analyzed in each experiment. Time-lapse images were acquired before the laser microirradiation. A 100% intensity scan (0.25 s) from a 405 nm laser was used to induce localized DNA damage in this study. To visualize green fluorescence, cells were exposed to 488 nm light illumination. The green fluorescence intensity at the irradiated sites was converted into numerical values using the SP8 software (Leica Microsystems). The relative green fluorescence intensity was calculated by subtracting the background intensity in the cells and dividing by the intensity in the irradiated area.

### Sodium dodecyl sulfate-polyacrylamide gel electrophoresis (SDS-PAGE) and western blotting

Cells were lysed in a radioimmunoprecipitation assay (RIPA) buffer (50 mM Tris HCl, pH 8.0, 250 mM NaCl, 25 mM ethylenediaminetetraacetic acid (EDTA), 0.5% v/v Triton X-100, 0.5% w/v SDS, and 0.5% w/v sodium deoxycholate) containing protease inhibitor cocktail (Nacalai Tesque Inc., cat# 25955-11) and phosphatase inhibitor cocktail (Nacalai Tesque Inc., cat# 07575-51), and the protein concentration was measured by a bicinchoninic acid (BCA) assay kit (Takara Bio) using bovine serum albumin as the standard. In all experiments, 20 μg of protein was loaded onto SDS-PAGE plates. The proteins were electrophoresed at 30 mA/gel plate for 1–1.5 h and transferred onto a polyvinylidene fluoride (PVDF) membrane at 100 V for 1.5 h. Next, the PVDF membrane was blocked with either 2% w/v BSA/TBS-T (tris-buffered saline and Tween 20) or 5% w/v skim milk/TBS-T for 1 h at room temperature on a shaker. For primary antibody reactions, the following primary antibodies were used for 1–4 h at room temperature: PNKP (rabbit, 1:1000, Novus, cat# NBP1-87257), PNKP (rabbit, 1:1000, Abcam, cat# ab181107), pT118-PNKP (rabbit, 1:1000, generated in this paper), pS114-PNKP (rabbit, 1:1000, generated in this paper), XRCC1 (mouse, 1:1000, Invitrogen, Thermo Fisher Scientific, cat# MA5-13412), XRCC4 (rabbit, 1:1000, generated in our laboratory ^60^), PCNA (rabbit, 1:500, Santa Cruz Biotechnology, cat# sc-7907), KAP1 (rabbit, 1:1000, abcam, cat# ab10483), pS824-KAP1 (rabbit, 1:1000, BETHYL, cat# A300-767A-2), Cyclin-A2 (mouse, 1:1000, Cell signaling, cat# BF683), Cyclin-E1 (rabbit, 1:1000, Sigma-Aldrich, cat# C4976), GFP (mouse, 1:3000, Nacali Tesque Inc., cat# GF200), ATM (mouse, 1:2000, Sigma Aldrich, cat# A1106), DNA-PKcs (rabbit, 1:2000, abcam, cat# Y393), p53 (mouse, 1:5000, Santa Cruz Biotechnology, cat# sc-126), pS15-p53 (mouse, 1:500, abcam, cat# ab223868), RPA2 (mouse, 1:3000, abcam, cat# ab2175), GAPDH (mouse, 1:10000, EMD Millipore, cat# MAB374), FLAG-M2 (mouse, 1:1000, Sigma-Aldrich, cat# A8592), RFP (rabbit, 1:1000, MBL, PM005), and FEN1 (mouse, 1:500, Santa Cruz, sc-28355). The PVDF membrane was washed three times with TBS-T. For secondary antibody reactions, horseradish peroxidase (HRP)-conjugated rabbit or mouse antibodies (Dako, cat# P0399 or P0447, respectively) were incubated for 1 h at room temperature. After washing five times with TBS-T, the membranes were developed using enhanced chemiluminescence (LI-COR Biosciences) and detected using a C-digit (LI-COR, Biosciences).

### Immunoprecipitation (IP)

For sample preparation for IP, HEK293 cells were grown on 100-mm dishes, washed twice in phosphate-buffered saline (PBS; Nacalai Tesque Inc.), and lysed in lysis buffer (50 mM Tris-HCl, pH 7.5, 100 mM NaCl, 0.2% NP-40, 1 mM MgCl_2_, and 10% glycerol) supplemented with cocktails of protease inhibitors and phosphatase inhibitors. After incubation for 30 min on the rotator at 4 °C, lysates were cleared by centrifugation 20,000 ×*g* for 20 mins at 4 °C. Next, lysates were incubated with 10 μL of GFP-Trap magnetic agarose beads (ChromoTek, GmbH) for 4 h with mixing on a rotator at 4 °C. The beads were then washed five times with lysis buffer, and proteins were eluted in 2 × SDS sample buffer (125 mM Tris-HCl, pH 6.8, 4% w/v SDS, 20% v/v glycerol, 0.01% w/v bromophenol blue, and 5% v/v 2-mercaptoethanol).

### Immunofluorescence

Cells were grown on glass coverslips and fixed with 4% formaldehyde for 15 min at 4 °C. Cells were subsequently permeabilized with PBS containing 0.2% Triton X-100 for 5 min at 4 °C. Following 30 min of blocking in PBS supplemented with 2% w/v BSA, primary antibody reactions were performed in PBS-T supplemented with 1% w/v BSA for 2 h at room temperature. Cells were washed three times with PBS, and secondary antibody reactions were performed in PBS-T supplemented with 1% BSA for 1 h at room temperature in the dark. After washing five times with PBS, coverslips were mounted in mounting medium (Dako) containing the nuclear staining dye 4’,6-diamidino-2-phenylindole dihydrochloride (DAPI) and allowed to dry for 2 h at room temperature in the dark. For the primary antibody, an anti-γH2AX mouse antibody (Merck Millipore, JBW301) was used at a 1:1000 dilution. An Alexa Fluor 594-conjugated mouse secondary antibody (Invitrogen, Thermo Fisher Scientific, cat# A32742) was used at a 1:2000 dilution.

To quantify γH2AX foci formation, nuclei and foci-positive cells were counted using the ImageJ software. Foci-positive cells were defined as those containing > 10 foci, and at least 100 cells were counted. Representative images are presented.

To measure ADP-ribosylation, cells were pretreated with 10 μM poly (ADP-ribose) glycohydrolase inhibitor (PARGi, TOCRIS Bio-Techne, PDD 00017273) for 30 min prior to IR exposure to increase total ADP-ribosylation levels by inhibiting hydrolysis of the ribose-ribose bonds present in poly (ADP-ribose). For the HU and EME assays, cells were incubated in 2 mM HU (Sigma-Aldrich) for 2 h or in 1 μM EME (Bio vision, BVN-B2339-50-50) for 1 h, with PARG inhibitor added during the final 20 min. As the primary reaction to detect SSBs, a PAN ADP-ribose binding reagent (rabbit, Merck, cat# 9QQ12P) was used at 1:1000 dilution in PBS-T supplemented with 1% w/v BSA. Alexa Fluor 488- or 594-conjugated rabbit secondary antibodies (Invitrogen, cat# A32731 or A32740, respectively) was used at a 1:2000 dilution. The mean intensity of ADP-ribose in nuclei stained with DAPI was measured using ImageJ software. At least 100 cells were counted, and the average ADP-ribose intensity was calculated using GraphPad Prism 8 (GraphPad Software Inc.).

To measure genome instability, cells with micronuclei and chromosome bridges were counted using DAPI staining. At least 300 cells were counted using ImageJ software.

### γ-ray irradiation

To examine the sensitivity and response to IR, cells were irradiated using ^60^Co γ-ray source in Tokyo Institute of Technology. The dose rate was measured using an ionizing chamber-type exposure dosimeter C-110 (Oyo Giken, Tokyo, Japan) and corrected for decay.

### Colony formation assay

The surviving fraction was determined using the colony formation assay. The cells were plated on 100 mm dishes. The number of plated cells was adjusted using higher doses of the indicated DNA-damaging agents to obtain an appropriate number of colonies. After incubation for 12–14 h at 37 °C under 5% CO_2_ conditions, cells were exposed to grading doses of γ-ray (1, 3, and 5 Gy), H_2_O_2_ (100 and 200 μM for 2 h), HU (1, 2, and 4 mM for 24 h) and CPT (Sigma-Aldrich) (5, 10, and 20 nM for 24 h). The cells were further incubated for 10–14 days to form colonies. After washing with PBS, cells were fixed with 99.5% ethanol and stained with staining solution (0.02% w/v crystal violet; 2.5% v/v methanol). After washing the plates twice with water and drying overnight, colonies containing more than 50 cells were counted manually. The plating efficiency was calculated as the number of colonies divided by the number of plated cells. The surviving fraction was calculated as the plating efficiency of the irradiated cells divided by the plating efficiency of the unirradiated cells. Experiments were independently repeated at least three times.

### Cell growth assay

Cell growth and growth rates were analyzed based on the number of cells at several time points ( Fig. 1F: days 0, 1, 2, 3, and 4; Fig. 2C and 2F: days 0, 1, and 4). Cells were prepared at 70– 80% confluency on 100-mm dishes or 60-mm dishes and spread onto 6-well plates (2 × 10^5^ cells/well). The cells were cultured at 37 °C in humidified atmosphere containing 5% CO_2_ conditions. After the indicated incubation period, cells were harvested by trypsinization, and cell numbers were counted using a Coulter counter (Beckman Coulter) in all experiments.

### Cell cycle distribution analysis by flowcytometry

The procedure for cell cycle distribution analysis has been described in our recent publication ^61^ and was appropriately modified for this study. In brief, the nascently synthesized DNA was labeled with EdU and Alexa Fluor 488 azide through Click reaction using a Click-iT EdU Imaging kit (Life Technologies, cat# C10337), and cells were then stained with propidium iodide (PI) using a Cell Cycle Phase Determination Kit (Cayman Chemicals, cat# 10009349) according to the manufacturer’s instruction. Cells grown in 6-well plates or 60-mm dish at 70–90% confluency were treated with 10 μM of EdU for 1 h and harvested by trypsinization. Harvested cells were washed with 0.1% w/v BSA/PBS and fixed/permeabilized by BD cytofix/cytoperm buffer (BD Biosciences, cat# 554714) according to the manufacturer’s instructions. Subsequently, the cells were washed with 1x BD perm/wash buffer and resuspended in a click-it reaction cocktail for 1 h at room temperature in the dark. After Click reaction, cells were washed with 1× BD perm/wash buffer and resuspend in PBS containing 0.02% w/v sodium azide, 0.02% w/v RNaseA, and 0.01% w/v PI for 1 h at room temperature in the dark. The cell suspension was supplemented with 500 mL of 0.1% w/v BSA/PBS for adjustment of the volume prior to the analysis and subjected to flow cytometry using Cell Lab Quanta SC (Beckman Coulter).

### DNA fiber analysis

The DNA fiber assay was performed according to a previously reported paper ^62^ and appropriately modified for this study. Cells grown on 60-mm dish at 70–90% confluency were initially labeled with 50 μM IdU for 15 min and subsequently labeled with 250 μM CldU for 15 min in a humidified CO_2_ incubator. Labeled cells were harvested by trypsinization and resuspended in ice-cold PBS at 1 × 10^6^–1 × 10^7^ cells/mL.

For the S1 nuclease assay, cells were labeled with 250 μM of CldU for 60 min with 10 μM of FEN1 inhibitor (MedChemExpress, FEN1-IN-3) and/or treated with 10 μM of PARP inhibitor (AdooQ Bioscience, Olaparib, AZD2281) in a humidified CO_2_ incubator, washed once with PBS, and permeabilized with CSK-100 buffer (100 mM NaCl, 10 mM HEPES pH7.8, 3 mM MgCl_2_, 300 mM sucrose, and 0.5% Triton X-100) for 10 min at RT. After permeabilization, cells were washed twice with ice-cold PBS and harvested by scraping. Harvested cells were divided into two 1.5 mL tubes for S1 nuclease treatment and subsequently centrifuged at 400 ×*g* for 3 min. Cell pellets were treated with S1 nuclease (Takara Bio, 2410A) for 30 min at 37 °C and resuspended in ice-cold PBS at 1 x 10^6^–1 x 10^7^ cells/mL.

2 μL of the cell suspension was spotted on one end of the glass slides (Matsunami glass, cat# S8215) and air-dried for 5 min. 7 mL of DNA fiber lysis buffer (200 mM Tris-HCl, pH 7.5, 50 mM EDTA, and 0.5% w/v SDS) were added to the cell suspension, gently stirred with a pipette tip, and incubated for 2 min. The glass slides were tilted at 15° to allow the fibers to spread along the slide and air-dried once the fiber solution reached the end of the glass slide. Glass slides were immersed in fixative (75% v/v methanol and 25% v/v acetic acid) and incubated for 10 min. After washing with distilled water twice, glass slides were immersed in 2.5 M HCl for 80 min, followed by three times wash with PBS, and blocked with 5% w/v BSA/PBS for 30 min. For primary antibody reaction, anti-BrdU (mouse, 1/100, BD biosciences, cat# 347580, reacts with IdU) and anti-BrdU (rat, 1/400, abcam, cat# ab6326, reacts with CldU) diluted in 5% w/v BSA/PBS were used and incubated in a humidified case for 2 h. After washing with PBS three times, goat anti-rat Alexa Fluor 488 (Invitrogen, 1/1000, cat# A110060) and goat anti-mouse Alexa Fluor 594 (Invitrogen, 1/1000, cat# A11005) were put onto the glass slides for secondary antibody reaction and incubated for 1 h in the dark. The glass slides were subsequently washed three times with PBS-T and mounted using mounting medium (Dako). To observe DNA fibers, an OLYMPUS IX71 (OLYMPUS) or Zeiss LSM880 (Carl Zeiss) fluorescence microscope was used, and at least 50 fibers were measured in each experiment. The tract length of the DNA fibers was measured using the ImageJ software and analyzed using GraphPad Prism 8 (GraphPad Software Inc.).

### Protein-DNA binding assay

An EpiQuik Colorimetric General Protein-DNA Binding Assay Kit (Epigentek Inc., cat# P-2004-96) was used to measure the DNA-binding ability of WT, T118A, and T118D PNKP, according to the manufacturer’s instructions. Nuclear extracts were harvested from U2OS *PNKP^−/−^* cells transfected with WT GFP-PNKP, T118A, or T118D expression vectors in non-denaturing lysis buffer (150 mM KCl, 50 mM Tris HCl, pH 8.3, 1 mM EDTA, and 1 mM DTT) supplemented with protease inhibitors. To measure the DNA binding ability to the gapped DNA, a biotinylated oligonucleotide (BioF20:5’-Biotin/TAGCACCTACCGATTGTATG/Phos-3’) and a non-biotinylated oligonucleotide (F15:5’/TACGTTTTTGTGTCG/3’) were annealed to a complementary strand oligonucleotide (R36:5’- Phos/CGACACAAAAACGTATCATACAATCGGTAGGTGCTA/3’) in annealing buffer (10 mM Tris, pH 7.5, 50 mM NaCl, and 1 mM EDTA). Oligonucleotides in annealing buffer were incubated at 95℃ for 5 min and cooled down slowly. 20 ng of biotinylated double-stranded oligonucleotides and 10 μg of nuclear extract were used for the DNA-binding reaction in streptavidin-coated tubes. Additionally, 1 μg/mL GFP antibody (Nacali Tesque Inc., cat# GF200) and 0.5 μg/mL HRP-conjugated mouse antibody (Dako, cat# P0447) were used to detect DNA-binding proteins. The absorbance was measured at 450 nm using an iMark Microplate Absorbance Reader (Bio-Rad Laboratories).

### PNKP phosphatase and kinase activity biochemical assay

The phosphatase and kinase activities of PNKP were determined in accordance with previous studies ^15, 63^. Fluorescently-labeled oligonucleotides (Integrated DNA Technologies) were used as PNKP substrates. For 3’-phosphatase reaction, ‘S1’ [5’ -(TAMRA) TAGCATCGATCAGTCCTC-3’ -P] and ‘C2’ [5’ -P-GAGGTCTAGCATCGTTAGTCA-(6-FAM)-3’] were annealed to a complementary strand oligonucleotide ‘B1’ [5’ - TGACTAACGATGCTAGACCTCTGAGGACTGATCGATGCTA-3’] in annealing buffer (10 mM Tris pH 7.5, 200 mM NaCl, and 1 mM EDTA). For 5’-kinase reaction, ‘C1’ [5’ -(TAMRA)- TAGCATCGATCAGTCCTC-3’ -OH] and ‘S2’ [5’ -OH-GAGGTCTAGCATCGTTAGTCA-(6-FAM)-3’] were annealed to ‘B1’ oligonucleotides in annealing buffer. Each of these oligonucleotide mixtures was treated at 95 °C for 5 min and then left at room temperature for 1 h to anneal and form substrate oligonucleotides. The expression vectors for WT PNKP, T118A, D171A, and K378A were transfected into *PNKP^-/-^*cells. In addition to these transfected *PNKP^−/−^* cells, WT U2OS and *PNKP^−/−^* cells were suspended in lysis buffer (25 mM Tris, pH 7.5, 10 mM EDTA, 10 mM EGTA, 100 mM NaCl, and 1% Triton X-100) and incubated for 15 min at 4 °C and then centrifuged at 16,000 ×*g* for 20 min at 4 °C. The supernatant was used as the cell-free protein extract. Protein extract of 1 × 10^5^ cells was incubated with 100 nM substrate oligonucleotides and 20 μM single-stranded nuclease competitor oligonucleotide [5’ - AAAGATCACAAGCATAAAGAGACAGG-3’] in reaction buffer (25 mM Tris, pH 7.5, 130 mM KCl, 10 mM MgCl2, 1 mM DTT, and 1 mM ATP) for 10 min at 37 °C. The enzymatic reactions were terminated by adding 25 μL of quenching buffer (90% formamide, 50 mM EDTA, 0.006% Orange G) to 25 μL of reaction solution. Each reaction sample was diluted with quenching buffer (× 10). Subsequently, 10 μL of each reaction sample was separated on a 20% denaturing polyacrylamide gel (7 M Urea and TBE buffer) for 16 h (500 V, 10 mA) and analyzed on a Typhoon 9500 (GE Healthcare Life Science).

### Isolation of proteins on nascent DNA (iPOND)

iPOND experiments were performed according to a previous protocol paper ^64^ and appropriately modified for this study. HEK293 cells grown in 15 cm dishes were treated with 10 μM EdU for 10 min. After EdU labeling, cells were fixed in 10 mL of 1% formaldehyde/PBS on the dishes for 20 min at room temperature and then quenched by adding 1 mL of 1.25M glycine. Cells were harvested by scraping 5 min after quenching and washed three times with PBS. Cells were subsequently permeabilized with PBS containing 0.25% Triton X-100 for 30 min at room temperature and washed twice with PBS. Each sample was divided into two 1.5 mL tubes for a click reaction (with biotin-azide) and a no-click control (without biotin-azide). Click-iT reaction buffer (Thermo Fisher, cat#: C10269) and biotin-azide (Cayman Chemical, cat#: 13040) were used for the Click reaction, according to the manufacturer’s protocol. Cells were washed twice in PBS and subsequently lysed in iPOND lysis buffer (1% SDS and 50mM Tris HCl pH8.0) containing protease inhibitor cocktail. Samples were sonicated using a BRANSON 150 sonicator, centrifuged at 20,000 ×*g* for 10 min at room temperature, and diluted in a 1:1 volume of PBS containing a protease inhibitor cocktail. 20 μL (per sample) of Streptavidin-Magnetic beads (Thermofisher, cat#: 88816) were washed twice with iPOND lysis buffer and incubated with samples overnight at 4 °C in the dark. Bead-sample mixtures were washed once in iPOND lysis buffer, once with low salt buffer (1% Triton-X100, 2 mM Tris HCl pH8.0, 2 mM EDTA, and 150 mM NaCl), once with high salt buffer (1% Triton-X100, 2 mM Tris HCl pH8.0, 2 mM EDTA, and 500 mM NaCl), and once with iPOND lysis buffer. Proteins were eluted in 2× SDS sample buffer by incubating for 25 min at 95 °C. Samples were resolved on SDS-PAGE, and proteins were detected by immunoblotting.

### In vitro kinase assay

Purified human recombinant 6× His-tagged PNKP was generously provided by Dr. Michael Weinfeld. Purified human recombinant GST-tagged CDK4/Cyclin D1 (cat# PV4400), CDK2/Cyclin E1 (cat# PV6295), CDK1/Cyclin A2 (cat# PV6280), and 6× His-tagged CDK2/Cyclin A2 (cat# PV3267) were purchased from Thermo Fisher Scientific. For kinase reaction, 200 ng of each CDK/Cyclin complex was incubated with 1 μg of His-PNKP in kinase reaction buffer (25 mM Tris-HCl at pH 7.5, 5 mM MgCl_2_, 5 mM sodium pyrophosphate, 2 mM ATP, and 2 mM DTT) for 1 h at 30 ℃. Reactions were quenched by adding 2× SDS sample buffer and boiling for 10 mins at 98 ℃. Samples were processed for western blotting and detected using specific antibodies.

### Statistical analysis

Statistical analysis was performed using either GraphPad Prism 8 (GraphPad Software Inc.) or Microsoft Excel. Unpaired (two-tailed) t-tests were used to analyze the statistical significance of differences between the two experimental groups. One-way analysis of variance (ANOVA) followed by post-hoc tests were used to analyze the statistical significance between multiple experimental groups. Sample scales are indicated in the figure legends. All experiments were independently performed at least three times with similar results. In all experiments, no statistical significance (ns) is defined as p > 0.05, * denotes 0.01 < p ≦ 0.05, ** denotes 0.005 < p ≦ 0.01, *** denotes 0.001 < p ≦ 0.005, and **** denotes 0.0005 < p ≦ 0.001.

## Acknowledgments

Authors thank Mr. Isao Yoda at Co^60^ radiation center, Drs. Kimitoshi Denda, Hiromi Yanagihara, Hirofumi Nakano, and Daisuke Morishita for technical assistance, and Matsumoto laboratory member for critical discussion. Authors also thank to Dr. Micheal weinfeld to provide purified PNKP peptide. This work was supported by The Uehara Memorial Foundation [to MS], Takeda Science Foundation [to MS], Kato Memorial Bioscience Foundation [to MS], Japan Atomic Energy Agency [to MS], and Chubu Electric Power [to MS], Tokyo Tech Academy for Co-creative Education of Environment and Energy Science [to KT], Tokyo Tech Academy for Leadership [to KT], Grant-in-Aid for Scientific Research from Japan Society for the Promotion of Science [Grant Numbers JP22K12369 to MS, JP15H02817, JP17K20042, JP20H04334 to YM and JP18K11642 to MI], Grant-in-Aid for Japan Society for the Promotion of Science Fellows [Grant Number JP20J13601 to KT], Japan Society for the Promotion of Science Overseas Research Fellowships [to KT] and Radiation Effects Association [to MI].

## Author contributions

K. T. and M. S. designed this study. K.T. constructed plasmid DNA vectors. K. T., R. I., K. S., M. S., N. K., T. M., and M. S. performed cell biology experiments. K.T. and M.I. carried out laser micro-irradiation assays. K.T., Y.M. and M.S. purified PNKP phosphorylation-specific antibodies. T. M. performed biochemical assay. M. I., Y, M., and M. S. supervised and provided advice. K. T. and M. S. wrote the manuscript with comments from the authors.

## Data and materials availability

All data, code, and materials used in the analyses must be available in some form to any researcher for purposes of reproducing or extending the analyses. Include a note explaining any restrictions on materials, such as materials transfer agreements (MTAs). Include accession numbers to any data relevant to the paper and deposited in a public database; include a brief description of the dataset or model with the number. If all data are in the paper and supplementary materials, include the sentence, “All data are available in the main text or the supplementary materials.”

## Supplementary figure legends

**Fig. S1.**
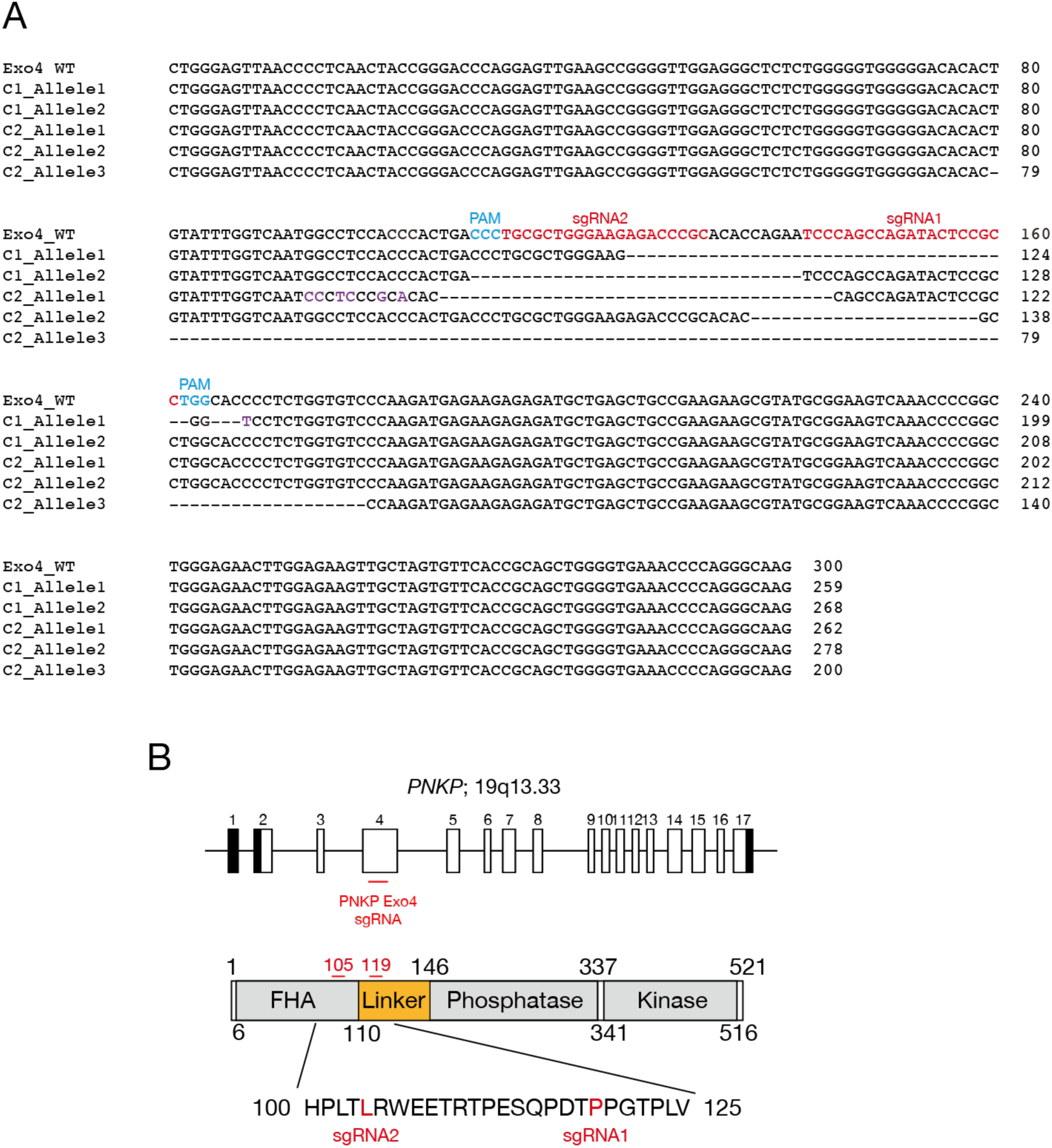
Generation of PNKP knockout U2OS cells by genome editing. A: DNA sequencing results in *PNKP* exon 4 of U2OS WT, *PNKP^−/−^*C1 and C2. DNA sequences of all alleles were aligned and PAM and sgRNA sequences were indicated. C1 and C2 have 2 and 3alleles, respectively, and all reading frames were frame-shifted. “Purple”, “Light blue” and “Red” characters indicated mutations, protospacer adjacent motif (PAM), single-guide RNA (sgRNA) sequences, respectively. “-” indicates a deletion of nucleotide. B: Targeting gene locus of CRISPR/Cas9 genome editing for generation of *PNKP^−/−^* cells. PNKP L105 and P119 located on exon 4 were targeted by Cas9 D10A.

**Fig. S2.**
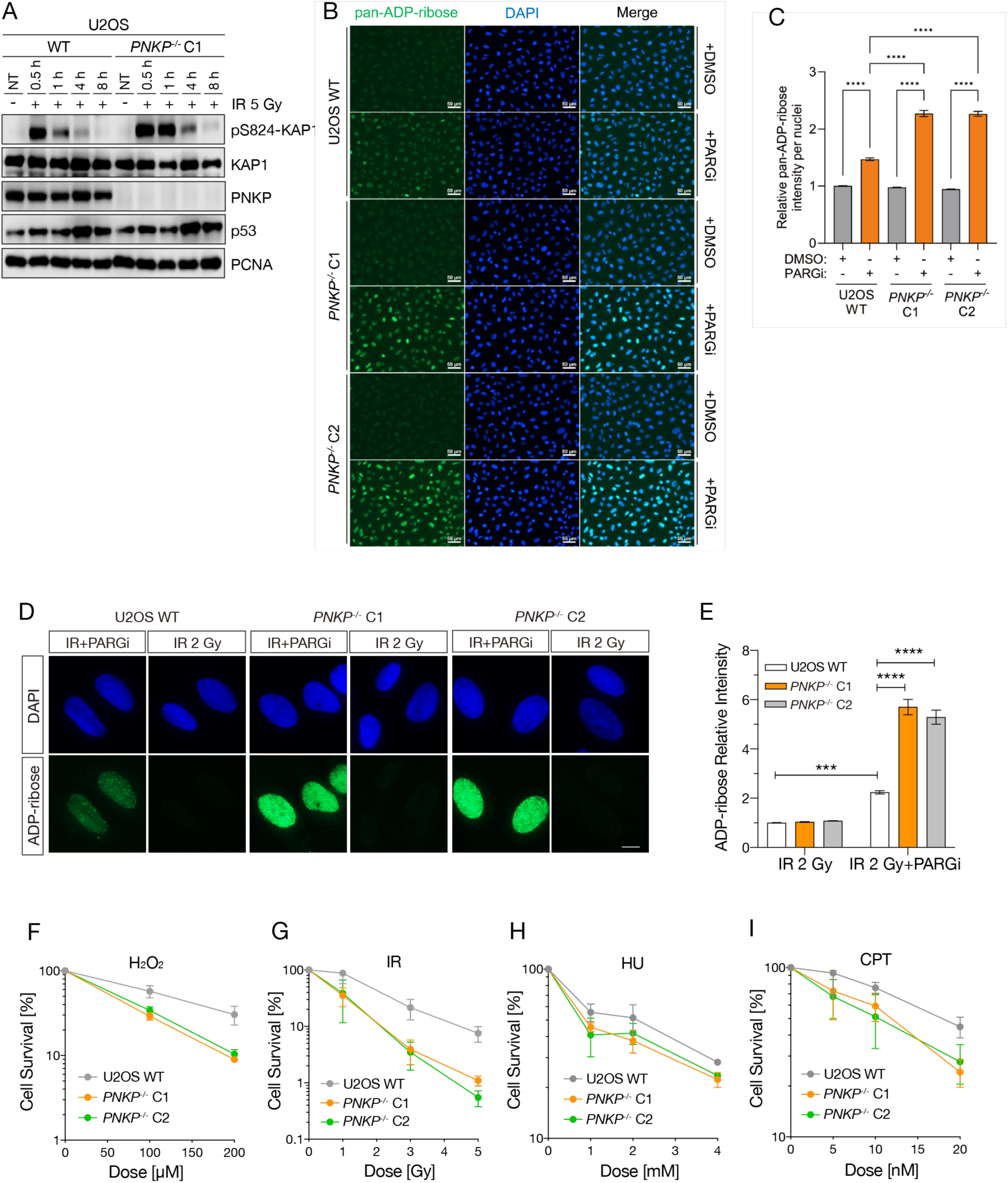
PNKP knockout cells showed high sensitivity to the genotoxic stress. A: DNA double-strand break repair activity in *PNKP^−/−^* C1 cells measured by western blotting. DNA double-strand breaks were analyzed by pS824-KAP1 antibody after indicated time points recovered from IR 5 Gy exposure. KAP1 and PCNA antibodies were used as loading control. p53 antibody was used as DNA damage response control. B and C: Measurement of endogenous DNA single-strand breaks of *PNKP^−/−^*cells. Single-strand breaks were analyzed by immunofluorescence using PAN-ADP-ribose binding reagents at 30min after IR 2 Gy exposure in U2OS WT and *PNKP^−/−^* (C1 and C2) cells treated with 10 μM PARGi for 60 min. D and E: Measurement of DNA single-strand break repair abilities of *PNKP^−/−^*cells. Single-strand breaks were analyzed by immunofluorescence using PAN-ADP-ribose binding reagents at 30min after IR 2 Gy exposure in U2OS WT and *PNKP^−/−^* (C1 and C2) cells treated with 10 μM PARGi for 30min prior to IR exposure. F and G: Cellular sensitivity of PNKP^−/−^ cells to the DNA damages, especially DNA strand breaks, was measured by colony formation assay. Cells were treated by hydrogen peroxide (H_2_O_2_: D) and IR (E) exposure at indicated dose. H and I: Cellular sensitivity of *PNKP^−/−^* cells to the DNA replication stress was measured by colony formation assay. Cells were treated by HU (F) and CPT (G) at indicated dose. In all panels, scale bar indicates 10 μm and error bars represent SEM. All experiments were performed at least three times independently. Statistical significance was assessed by one-way ANOVA with post hoc Sidak’s multiple comparisons test (\*\*\**P*<0.001, \*\*\*\**P*<0.0001).

**Fig. S3.**
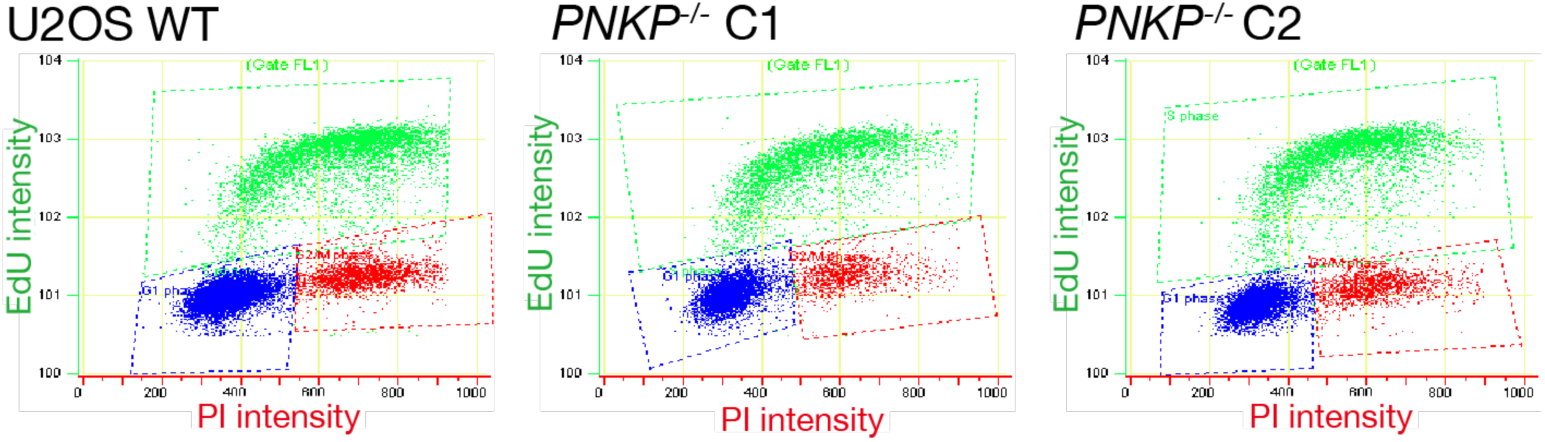
Cell cycle analysis in PNKP knockout cells s. Flowcytometric analysis of cell cycle distribution in U2OS WT, *PNKP^−/−^*C1 and C2 cells. Nascent synthesized DNA was detected by EdU (vertical axis) and whole DNA was stained by PI (horizontal axis). Each cell cycle was determined by gating of broken lines (Blue: G1 phase, Light green: S phase, Red: G2/M phase). Quantified result was shown in Fig. 1B.

**Fig. S4.**
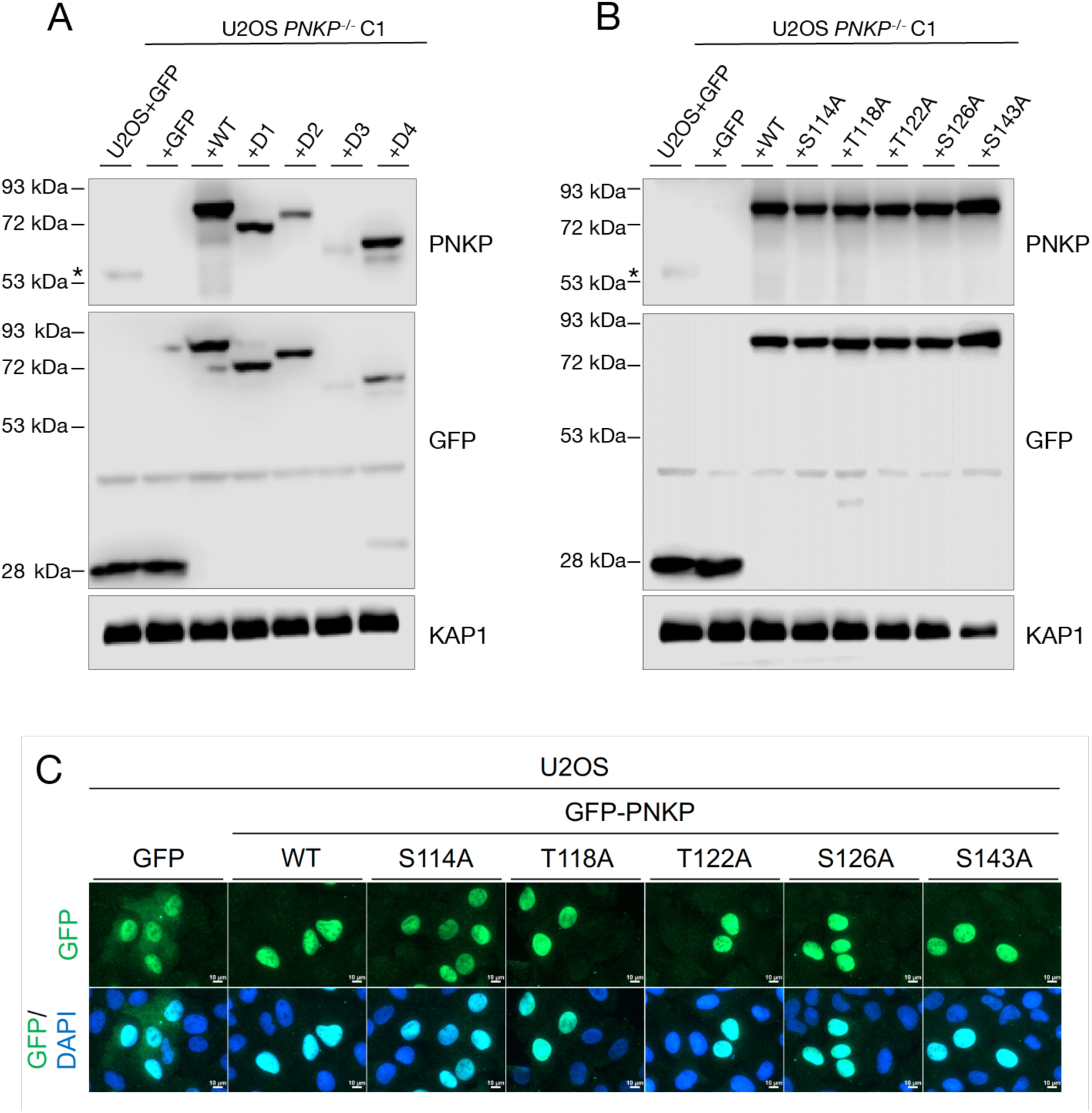
PNKP mutants expressing in knockout cells. A: Protein expression analysis of U2OS WT and *PNKP^−/−^* C1 cells transiently expressing GFP or indicated PNKP deletion mutants. Protein expression was detected by western blotting. GFP antibody was used for confirming the expression of exogenous PNKP deletion mutants. PNKP antibody was used for comparing expression levels of endogenous and exogenous PNKP. KAP1 antibody was used as a loading control. “*” indicates endogenous PNKP. B and C: Protein expression analysis of U2OS WT and *PNKP^−/−^* C1 cells transiently expressing GFP or indicated PNKP point mutants. Protein expression was detected by western blotting (B) and immunofluorescence (C). (B) GFP antibody was used for confirming the expression of exogenous PNKP point mutants. PNKP antibody was used for comparing expression levels of endogenous and exogenous PNKP. KAP1 antibody was used as a loading control. “*” indicates endogenous PNKP. (C) GFP fluorescence was observed by fluorescent microscope. DAPI was used as nucleus staining. In all panels, scale bar indicates 10 μm and error bars represent SEM.

**Fig. S5.**
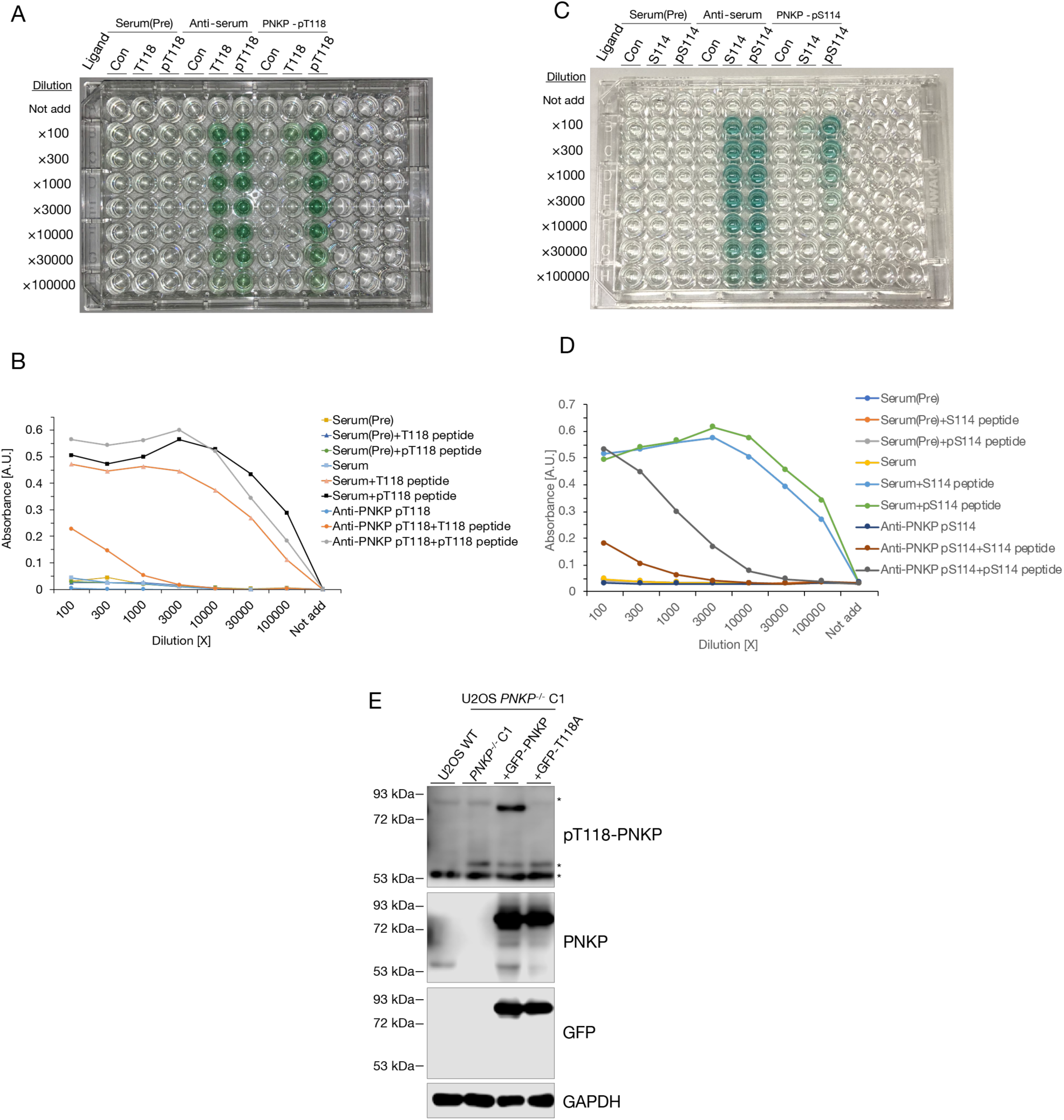
Generation of phosphorylation of T118 or S114 PNKP recognized antibodies. A: Representative image of ELISA assay for pT118-PNKP antibody. Dilutions were shown in vertical axis, and ligands and antibodies were shown in horizontal axis. B: Quantified results of Fig. S3A for confirming the titer and specificity of pT118-PNKP antibody. Absorbance was shown in vertical axis, and dilutions were shown in horizontal axis. C: Representative image of ELISA assay for pS114-PNKP antibody. Dilutions were shown in vertical axis, and ligands and antibodies were shown in horizontal axis. D: Quantified results of Fig. S3D for confirming the titer and specificity of pS114-PNKP antibody. Absorbance was shown in vertical axis, and dilutions were shown in horizontal axis. E: Specificity of pT118-PNKP antibody for cell lysate from U2OS WT and *PNKP^−/−^*C1 cells transiently expressing GFP-PNKP or GFP-PNKP T118A mutant. Asterisks indicate non-specific detections. GFP and PNKP antibodies were used for confirming the expression of GFP-PNKP and GFP-PNKP T118A mutant. GAPDH antibody was used as a loading control.

**Fig. S6.**
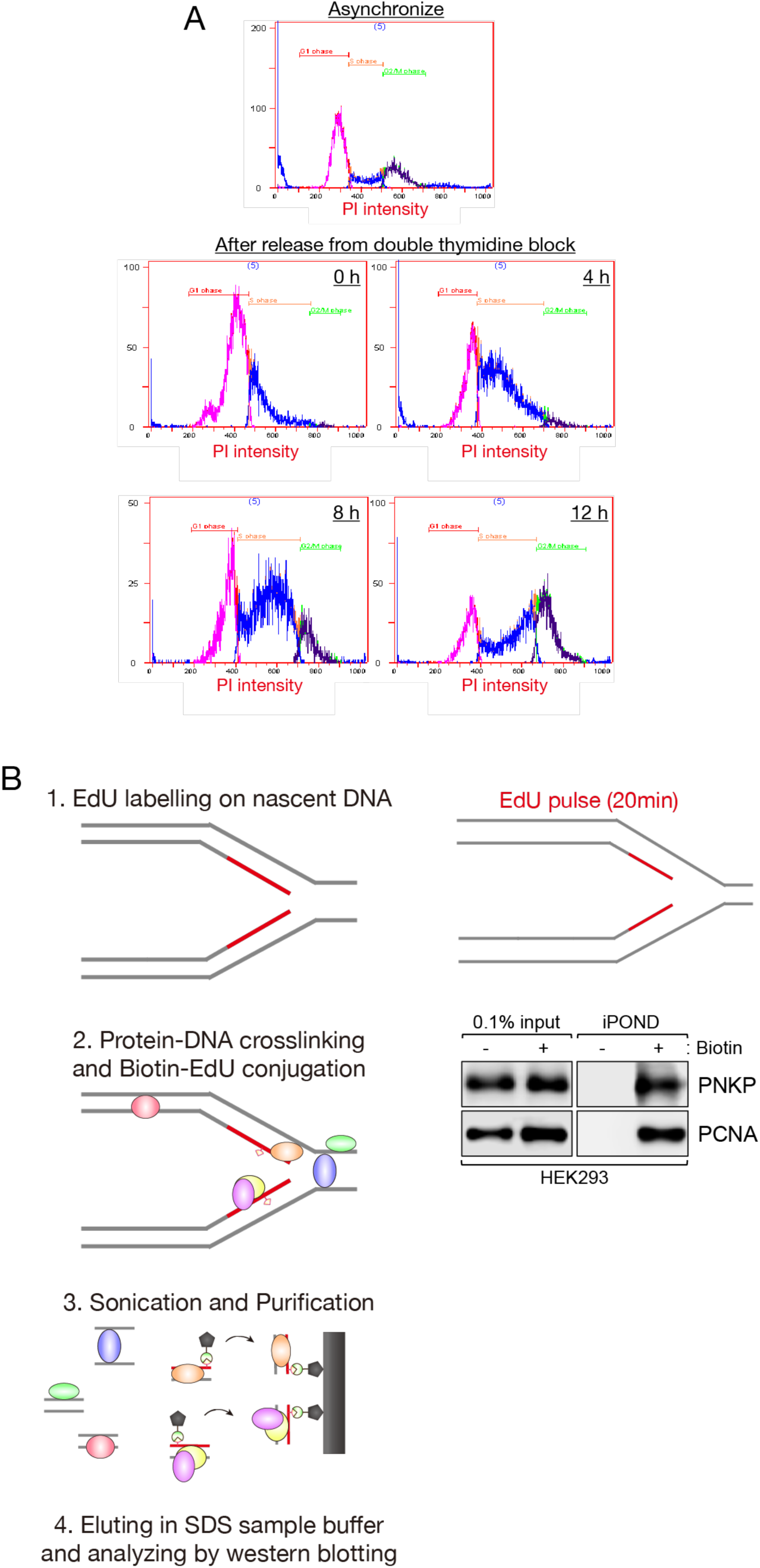
Cell cycle analysis of PNKP and scheme of iPOND. A: Flowcytometric analysis of cell cycle distribution of HCT116 cells transiently expressing GFP-PNKP at indicated time after release from double thymidine block. Cell cycle was determined by DNA contents measured by PI staining (horizontal axis). B: Schematic of iPOND experiment. Proteins bound to EdU labeled nascent DNA in HEK293 cells were isolated using click reaction with biotin-azide followed by streptavidin-pulldowns. 0.1% of lysate used in Streptavidin-pulldowns represented as 0.1% input as a loading control.

**Fig. S7.**
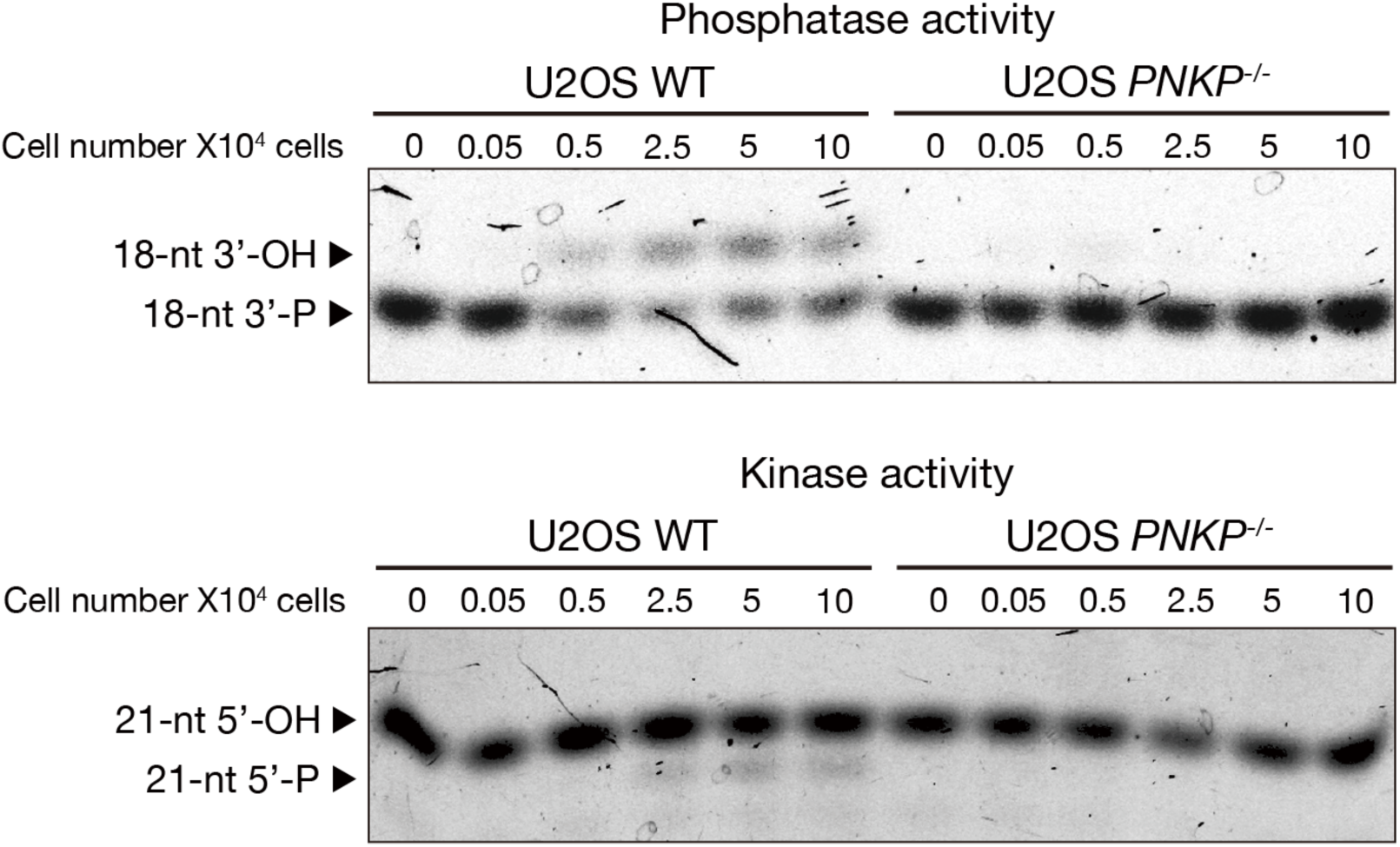
PNKP phosphatase and kinase activity biochemical analysis. Indicated number of cell extracts harvested from U2OS WT and *PNKP^−/−^* C1 cells were incubated with TAMRA or 6-FAM labelled oligonucleotide duplex harboring a SSB GAP structure. Arrows indicate the positions of the TAMRA labelled phosphatase substrates (top) and 6-FAM labelled kinase substrates (bottom).

**Fig. S8.**
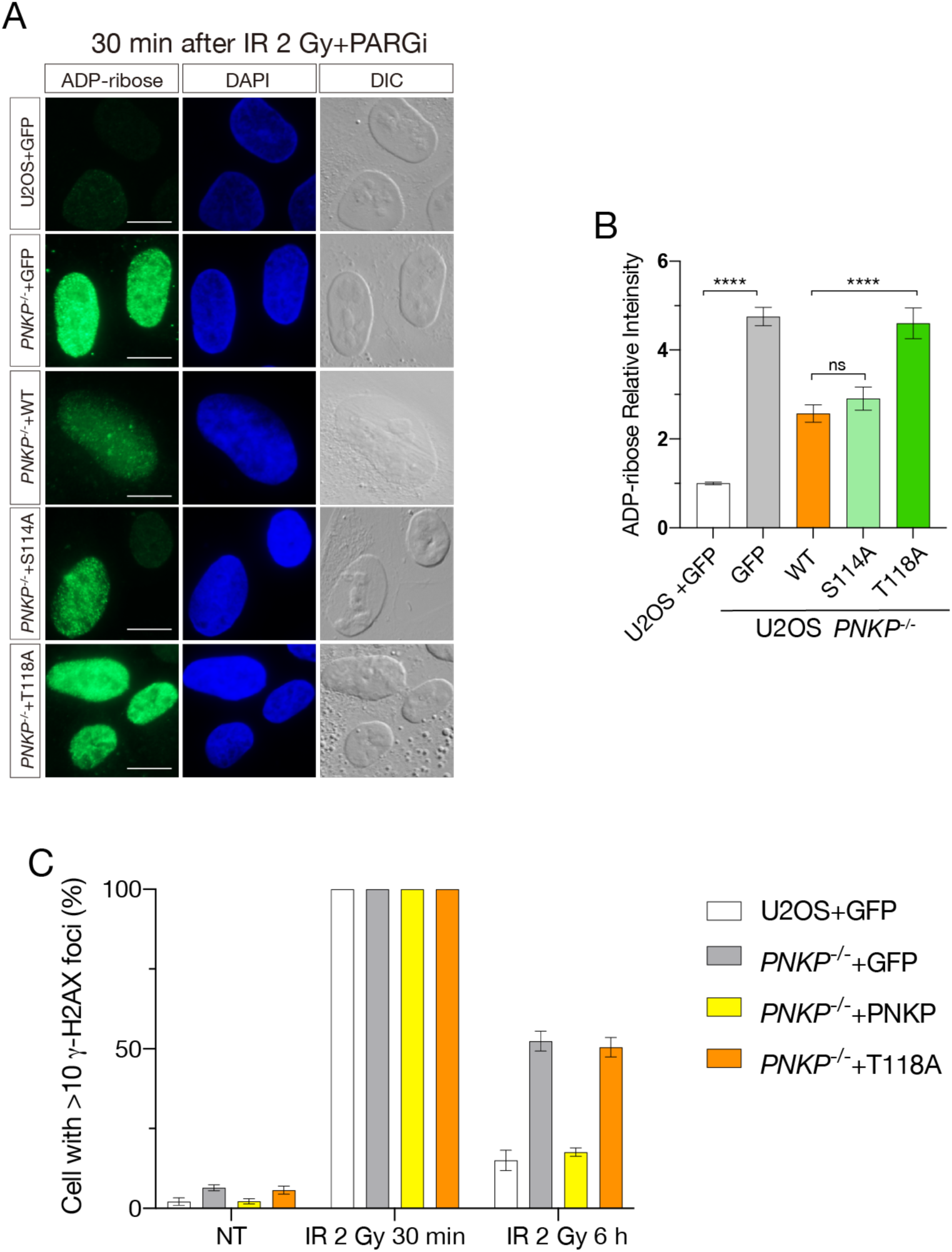
Phosphorylation of PNKP at T118 is required for genome stability. A: Representative images of immunofluorescence using PAN ADP-ribose binding reagents at 30 min after 2Gy IR exposure in U2OS WT and *PNKP^−/−^* cells transiently expressing GFP or indicated PNKP mutants. PARGi was added 30 min prior to IR exposure. Images detected by ADP-ribose, DAPI and DIC were shown. Scale bar indicates as 10 μm. B: Quantified result of Fig. S6A. Relative ADP-ribose intensity is shown in vertical axis and cell types were shown in horizontal axis. At least 100 cells were analyzed for the quantification. Statistical significance was indicated as not significant (ns) and ****: 0.0005 < p ≦ 0.001. C: Measurement of DNA DSB repair ability of indicated cells after 2 Gy IR exposure. Cells were harvested at 30 min and 6 h after IR exposure. Percentage of γH2AX positive cells is shown in vertical axis and conditions were shown in horizontal axis. NT indicates non-treatment. At least 100 cells were analyzed for the quantification. Statistical significance was indicated as not significant (ns) and ****: 0.0005 < p ≦ 0.001.

**Fig. S9.**
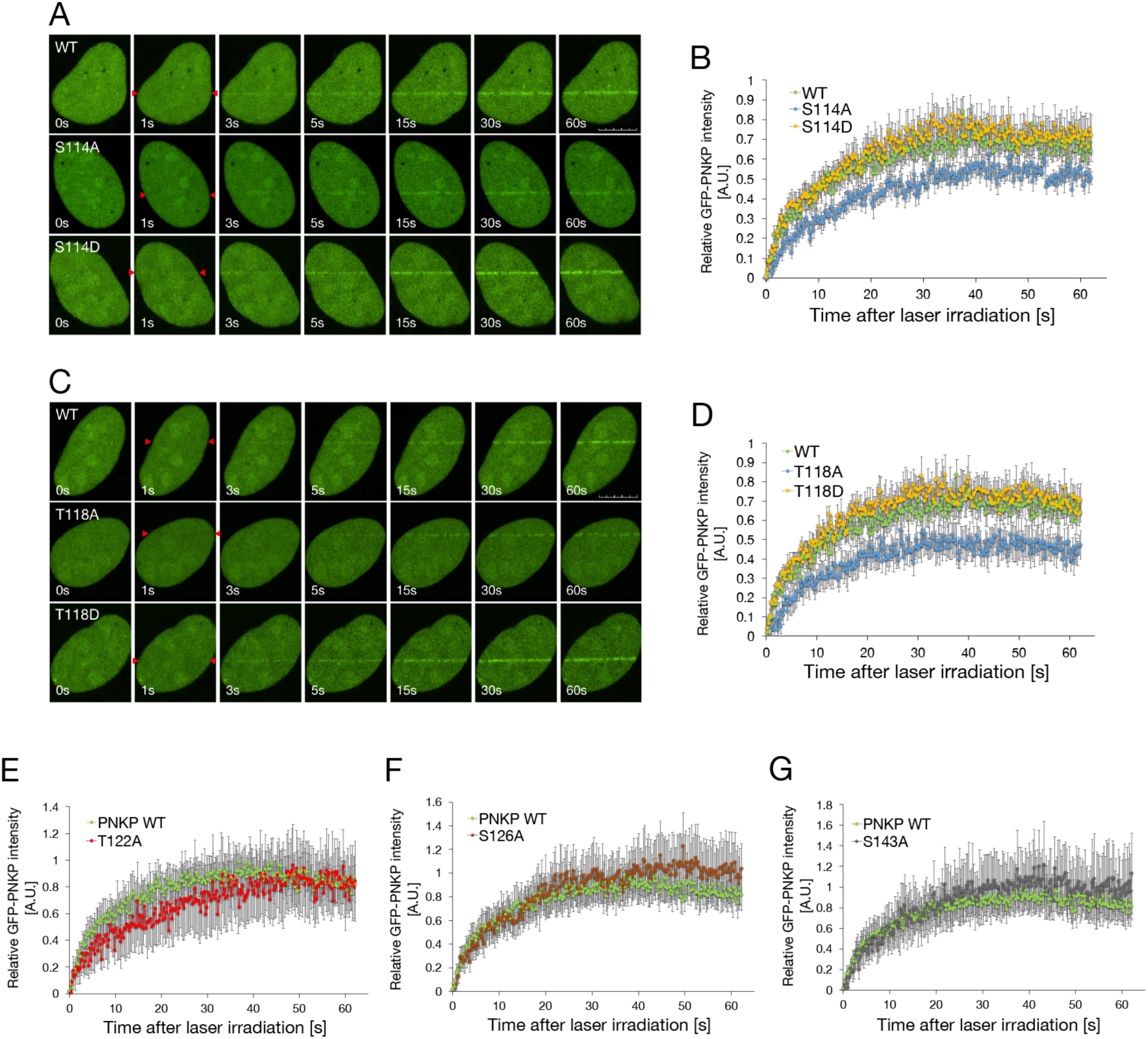
Linker region of PNKP is required for recruitment to the DNA damage sites. A : Representative live-cell images of the GFP-tagged PNKP S114 substituted mutants (S114A and S114D) after laser micro-irradiation. Micro-laser was irradiated on the line between red arrows. Cells were sensitized by Hoechst 33258 at 30 min prior to the laser micro-irradiation. Scale bar indicates as 10 μm. B: Relative green fluorescence intensity of indicated PNKP S114 substituted mutants after laser micro-irradiation. Green fluorescence intensity was scored every 0.25 s up to 60 s. 20 cells were scored at each point. Error bars represent SEM. C: Representative live-cell images of the GFP-tagged PNKP T118 substituted mutants (T118A and T118D) after laser micro-irradiation. Micro-laser was irradiated on the line between red arrows. Scale bar indicates as 10 μm. D: Relative green fluorescence intensity of indicated PNKP T118 substituted mutants (T118A and T118D) after laser micro-irradiation. 20 cells were scored at each point. Error bars represent SEM. E, F and G: Relative green fluorescence intensity of indicated PNKP point substituted mutants (T122A (E), S126A (F) and S143A (G)) after laser micro-irradiation. 20 cells were scored at each point. Error bars represent SEM.

**Supplementary table 1.**
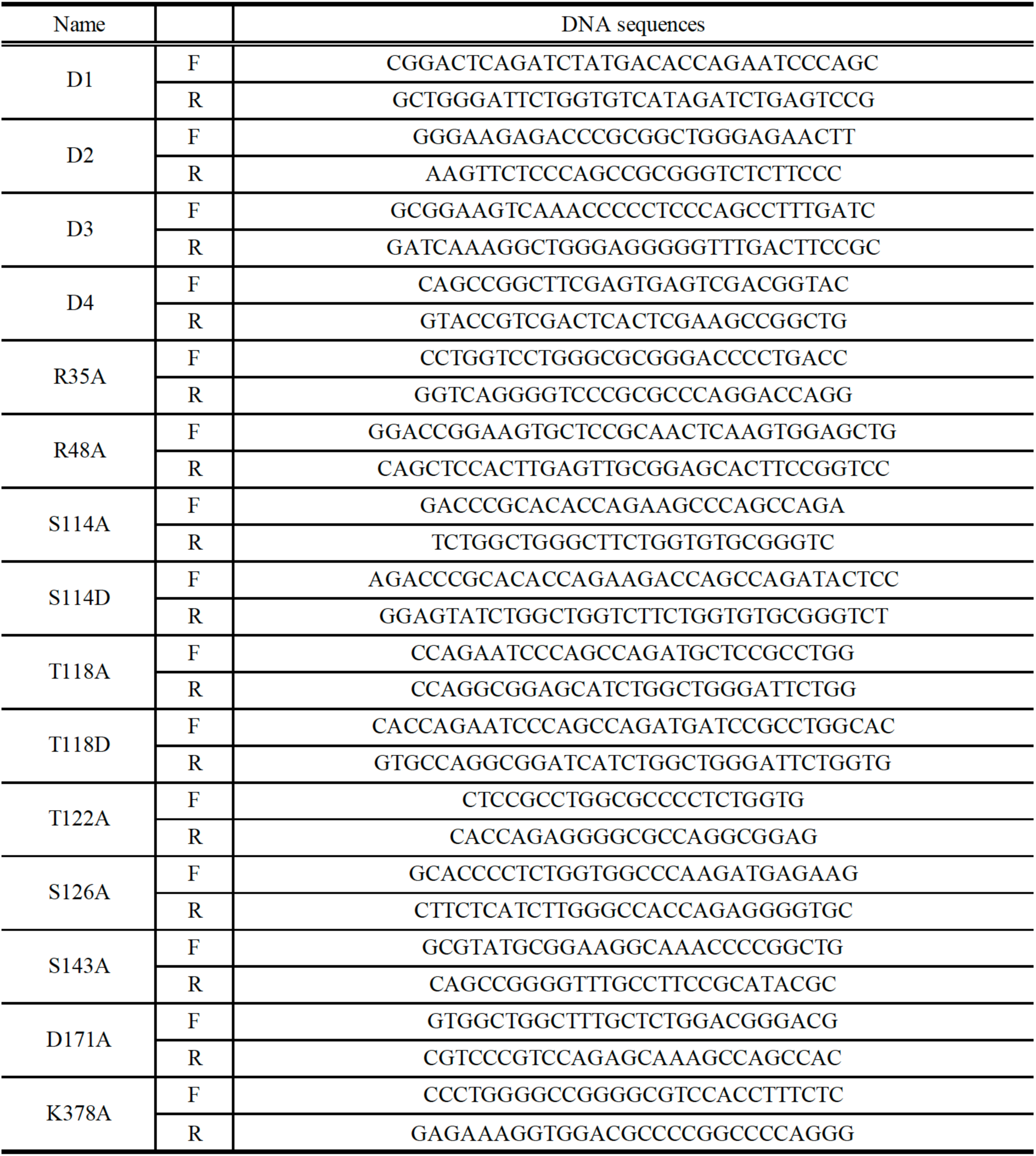
Sequences of DNA oligonucleotide primers for mutagenesis of PNKP. “F” and “R” indicate forward and reverse sequences, respectively.

**Supplementary table 2.**
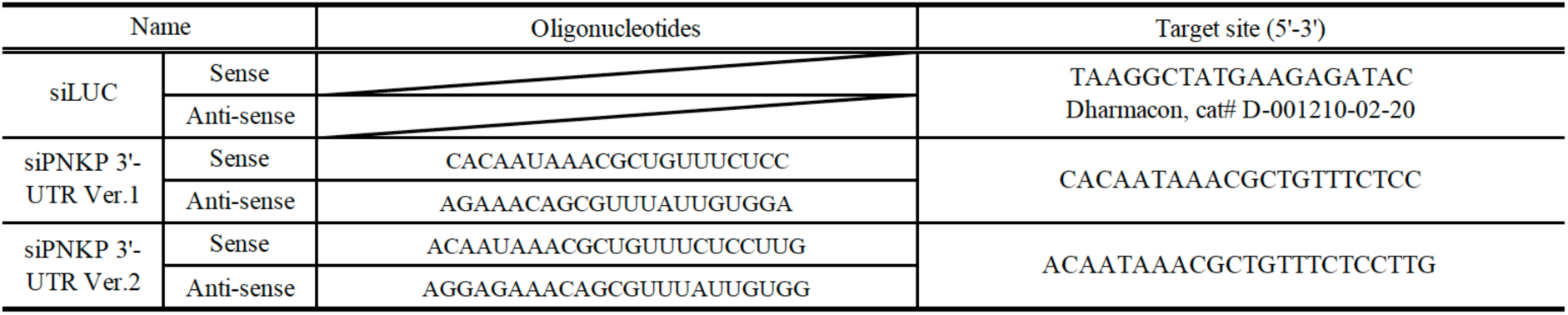
Oligonucleotide sequences of siRNAs for specific depletion with indicated proteins.

## References

1. Chappell C, Hanakahi LA, Karimi-Busheri F, Weinfeld M, West SC. Involvement of human polynucleotide kinase in double-strand break repair by non-homologous end joining. EMBO J 21, 2827–2832 (2002).

2. Karimi-Busheri F, et al. Molecular characterization of a human DNA kinase. J Biol Chem 274, 24187–24194 (1999).

3. Jilani A, et al. Molecular cloning of the human gene, PNKP, encoding a polynucleotide kinase 3’-phosphatase and evidence for its role in repair of DNA strand breaks caused by oxidative damage. J Biol Chem 274, 24176–24186 (1999).

4. Breslin C, Caldecott KW. DNA 3’-phosphatase activity is critical for rapid global rates of single-strand break repair following oxidative stress. Mol Cell Biol 29, 4653–4662 (2009).

5. Mani RS, et al. Domain analysis of PNKP-XRCC1 interactions: Influence of genetic variants of XRCC1. J Biol Chem 294, 520–530 (2019).

6. Mani RS, et al. Dual modes of interaction between XRCC4 and polynucleotide kinase/phosphatase: implications for nonhomologous end joining. J Biol Chem 285, 37619–37629 (2010).

7. Tsukada K, Matsumoto Y, Shimada M. Linker region is required for efficient nuclear localization of polynucleotide kinase phosphatase. PLoS One 15, e0239404 (2020).

8. Tsukada K, Shimada M, Imamura R, Saikawa K, Ishiai M, Matsumoto Y. The FHA domain of PNKP is essential for its recruitment to DNA damage sites and maintenance of genome stability. Mutat Res 822, 111727 (2020).

9. Segal-Raz H, et al. ATM-mediated phosphorylation of polynucleotide kinase/phosphatase is required for effective DNA double-strand break repair. EMBO Rep 12, 713–719 (2011).

10. Parsons JL, et al. Phosphorylation of PNKP by ATM prevents its proteasomal degradation and enhances resistance to oxidative stress. Nucleic Acids Res 40, 11404–11415 (2012).

11. Shen J, et al. Mutations in PNKP cause microcephaly, seizures and defects in DNA repair. Nat Genet 42, 245–249 (2010).

12. Bras J, et al. Mutations in PNKP cause recessive ataxia with oculomotor apraxia type 4. Am J Hum Genet 96, 474–479 (2015).

13. Pedroso JL, et al. Mutation in PNKP presenting initially as axonal Charcot-Marie-Tooth disease. Neurol Genet 1, e30 (2015).

14. Reynolds JJ, Walker AK, Gilmore EC, Walsh CA, Caldecott KW. Impact of PNKP mutations associated with microcephaly, seizures and developmental delay on enzyme activity and DNA strand break repair. Nucleic Acids Res 40, 6608–6619 (2012).

15. Kalasova I, et al. Pathological mutations in PNKP trigger defects in DNA single-strand break repair but not DNA double-strand break repair. Nucleic Acids Res, (2020).

16. Bermudez-Guzman L, Jimenez-Huezo G, Arguedas A, Leal A. Mutational survivorship bias: The case of PNKP. PLoS One 15, e0237682 (2020).

17. Siddiqui K, On KF, Diffley JF. Regulating DNA replication in eukarya. Cold Spring Harb Perspect Biol 5, (2013).

18. O’Donnell M, Langston L, Stillman B. Principles and concepts of DNA replication in bacteria, archaea, and eukarya. Cold Spring Harb Perspect Biol 5, (2013).

19. Leonard AC, Mechali M. DNA replication origins. Cold Spring Harb Perspect Biol 5, a010116 (2013).

20. Okazaki R, Okazaki T, Sakabe K, Sugimoto K, Sugino A. Mechanism of DNA chain growth. I. Possible discontinuity and unusual secondary structure of newly synthesized chains. Proc Natl Acad Sci U S A 59, 598–605 (1968).

21. Hanzlikova H, Kalasova I, Demin AA, Pennicott LE, Cihlarova Z, Caldecott KW. The Importance of Poly(ADP-Ribose) Polymerase as a Sensor of Unligated Okazaki Fragments during DNA Replication. Mol Cell 71, 319–331 e313 (2018).

22. Ray Chaudhuri A, et al. Topoisomerase I poisoning results in PARP-mediated replication fork reversal. Nat Struct Mol Biol 19, 417–423 (2012).

23. Ray Chaudhuri A, Nussenzweig A. The multifaceted roles of PARP1 in DNA repair and chromatin remodelling. Nat Rev Mol Cell Biol 18, 610–621 (2017).

24. Swaffer MP, Jones AW, Flynn HR, Snijders AP, Nurse P. CDK Substrate Phosphorylation and Ordering the Cell Cycle. Cell 167, 1750–1761 e1716 (2016).

25. Wold MS. Replication protein A: a heterotrimeric, single-stranded DNA-binding protein required for eukaryotic DNA metabolism. Annu Rev Biochem 66, 61–92 (1997).

26. Cortez D, Guntuku S, Qin J, Elledge SJ. ATR and ATRIP: partners in checkpoint signaling. Science 294, 1713–1716 (2001).

27. Zou L, Elledge SJ. Sensing DNA damage through ATRIP recognition of RPA-ssDNA complexes. Science 300, 1542–1548 (2003).

28. Shimada M, Dumitrache LC, Russell HR, McKinnon PJ. Polynucleotide kinase-phosphatase enables neurogenesis via multiple DNA repair pathways to maintain genome stability. EMBO J 34, 2465–2480 (2015).

29. Chiang TW, le Sage C, Larrieu D, Demir M, Jackson SP. CRISPR-Cas9(D10A) nickase-based genotypic and phenotypic screening to enhance genome editing. Sci Rep 6, 24356 (2016).

30. Hoch NC, et al. XRCC1 mutation is associated with PARP1 hyperactivation and cerebellar ataxia. Nature 541, 87–91 (2017).

31. Chalasani SL, et al. Persistent 3’-phosphate termini and increased cytotoxicity of radiomimetic DNA double-strand breaks in cells lacking polynucleotide kinase/phosphatase despite presence of an alternative 3’-phosphatase. DNA Repair (Amst) 68, 12–24 (2018).

32. Thakar T, et al. Ubiquitinated-PCNA protects replication forks from DNA2-mediated degradation by regulating Okazaki fragment maturation and chromatin assembly. Nat Commun 11, 2147 (2020).

33. Hornbeck PV, Zhang B, Murray B, Kornhauser JM, Latham V, Skrzypek E. PhosphoSitePlus, 2014: mutations, PTMs and recalibrations. Nucleic Acids Res 43, D512–520 (2015).

34. Zolner AE, et al. Phosphorylation of polynucleotide kinase/ phosphatase by DNA-dependent protein kinase and ataxia-telangiectasia mutated regulates its association with sites of DNA damage. Nucleic Acids Res 39, 9224–9237 (2011).

35. Fung TK, Ma HT, Poon RY. Specialized roles of the two mitotic cyclins in somatic cells: cyclin A as an activator of M phase-promoting factor. Mol Biol Cell 18, 1861–1873 (2007).

36. Pagano M, Pepperkok R, Verde F, Ansorge W, Draetta G. Cyclin A is required at two points in the human cell cycle. EMBO J 11, 961–971 (1992).

37. Honda R, et al. The structure of cyclin E1/CDK2: implications for CDK2 activation and CDK2-independent roles. EMBO J 24, 452–463 (2005).

38. Meijer L, et al. Biochemical and cellular effects of roscovitine, a potent and selective inhibitor of the cyclin-dependent kinases cdc2, cdk2 and cdk5. Eur J Biochem 243, 527–536 (1997).

39. Bukanov NO, Smith LA, Klinger KW, Ledbetter SR, Ibraghimov-Beskrovnaya O. Long-lasting arrest of murine polycystic kidney disease with CDK inhibitor roscovitine. Nature 444, 949–952 (2006).

40. Burhans WC, Vassilev LT, Wu J, Sogo JM, Nallaseth FS, DePamphilis ML. Emetine allows identification of origins of mammalian DNA replication by imbalanced DNA synthesis, not through conservative nucleosome segregation. EMBO J 10, 4351–4360 (1991).

41. Lukac D, Machacova Z, Moudry P. Emetine blocks DNA replication via proteosynthesis inhibition not by targeting Okazaki fragments. Life Sci Alliance 5, (2022).

42. Exell JC, et al. Cellularly active N-hydroxyurea FEN1 inhibitors block substrate entry to the active site. Nat Chem Biol 12, 815–821 (2016).

43. Ward TA, McHugh PJ, Durant ST. Small molecule inhibitors uncover synthetic genetic interactions of human flap endonuclease 1 (FEN1) with DNA damage response genes. PLoS One 12, e0179278 (2017).

44. Zheng L, Shen B. Okazaki fragment maturation: nucleases take centre stage. J Mol Cell Biol 3, 23–30 (2011).

45. Kalasova I, et al. Novel PNKP mutations causing defective DNA strand break repair and PARP1 hyperactivity in MCSZ. Neurol Genet 5, e320 (2019).

46. Vaitsiankova A, et al. PARP inhibition impedes the maturation of nascent DNA strands during DNA replication. Nat Struct Mol Biol 29, 329–338 (2022).

47. Kolas NK, et al. Orchestration of the DNA-damage response by the RNF8 ubiquitin ligase. Science 318, 1637–1640 (2007).

48. Blackford AN, Jackson SP. ATM, ATR, and DNA-PK: The Trinity at the Heart of the DNA Damage Response. Mol Cell 66, 801–817 (2017).

49. Cortez D, Wang Y, Qin J, Elledge SJ. Requirement of ATM-dependent phosphorylation of brca1 in the DNA damage response to double-strand breaks. Science 286, 1162–1166 (1999).

50. Laney JD, Hochstrasser M. Substrate targeting in the ubiquitin system. Cell 97, 427–430 (1999).

51. Kunkel TA. DNA replication fidelity. J Biol Chem 279, 16895–16898 (2004).

52. Genois MM, et al. CARM1 regulates replication fork speed and stress response by stimulating PARP1. Mol Cell 81, 784–800 e788 (2021).

53. Maya-Mendoza A, Moudry P, Merchut-Maya JM, Lee M, Strauss R, Bartek J. High speed of fork progression induces DNA replication stress and genomic instability. Nature 559, 279–284 (2018).

54. Laipis PJ, Levine AJ. DNA replication in SV40-infected cells. IX. The inhibition of a gap-filling step during discontinuous synthesis of SV40 DNA. *Virology* **56**, 580-594 (1973).

55. Magnusson G. Hydroxyurea-induced accumulation of short fragments during polyoma DNA replication. II. Behavior during incubation of isolated nuclei. J Virol 12, 609–615 (1973).

56. Magnusson G. Hydroxyurea-induced accumulation of short fragments during polyoma DNA replication. I. Characterization of fragments. J Virol 12, 600–608 (1973).

57. Freschauf GK, et al. Identification of a small molecule inhibitor of the human DNA repair enzyme polynucleotide kinase/phosphatase. Cancer Res 69, 7739–7746 (2009).

58. Inano S, et al. RFWD3-Mediated Ubiquitination Promotes Timely Removal of Both RPA and RAD51 from DNA Damage Sites to Facilitate Homologous Recombination. Mol Cell 66, 622–634 e628 (2017).

59. Mochizuki AL, et al. PARI Regulates Stalled Replication Fork Processing To Maintain Genome Stability upon Replication Stress in Mice. Mol Cell Biol 37, (2017).

60. Kamdar RP, Matsumoto Y. Radiation-induced XRCC4 association with chromatin DNA analyzed by biochemical fractionation. J Radiat Res 51, 303–313 (2010).

61. Tsuchiya H, Shimada M, Tsukada K, Meng Q, Kobayashi J, Matsumoto Y. Diminished or inversed dose-rate effect on clonogenic ability in Ku-deficient rodent cells. J Radiat Res 62, 198–205 (2021).

62. Schwab RA, Niedzwiedz W. Visualization of DNA replication in the vertebrate model system DT40 using the DNA fiber technique. J Vis Exp, e3255 (2011).

63. Dobson CJ, Allinson SL. The phosphatase activity of mammalian polynucleotide kinase takes precedence over its kinase activity in repair of single strand breaks. Nucleic Acids Res 34, 2230–2237 (2006).

64. Dungrawala H, Cortez D. Purification of proteins on newly synthesized DNA using iPOND. Methods Mol Biol 1228, 123–131 (2015).

