## Supplementary Figure 1 for "CDKs-mediated phosphorylation of PNKP is required for end-processing of single-strand DNA gaps on Okazaki Fragments and genome stability"

A

```
                                                              PAM          sgRNA2                          sgRNA1
Exo4 WT       CTGGGAGTTAACCCCTCAACTACCGGGACCCAGGAGTTGAAGCCGGGGTTGGAGGGCTCTCTGGGGGTGGGGGACACACT  80
C1_Allele1    CTGGGAGTTAACCCCTCAACTACCGGGACCCAGGAGTTGAAGCCGGGGTTGGAGGGCTCTCTGGGGGTGGGGGACACACT  80
C1_Allele2    CTGGGAGTTAACCCCTCAACTACCGGGACCCAGGAGTTGAAGCCGGGGTTGGAGGGCTCTCTGGGGGTGGGGGACACACT  80
C2_Allele1    CTGGGAGTTAACCCCTCAACTACCGGGACCCAGGAGTTGAAGCCGGGGTTGGAGGGCTCTCTGGGGGTGGGGGACACACT  80
C2_Allele2    CTGGGAGTTAACCCCTCAACTACCGGGACCCAGGAGTTGAAGCCGGGGTTGGAGGGCTCTCTGGGGGTGGGGGACACACT  80
C2_Allele3    CTGGGAGTTAACCCCTCAACTACCGGGACCCAGGAGTTGAAGCCGGGGTTGGAGGGCTCTCTGGGGGTGGGGGACACAC-  79

                                             PAM          sgRNA2                          sgRNA1
Exo4_WT       GTATTTGGTCAATGGCCTCCACCCACTGACCCTGCGCTGGGAAGAGACCCGCACACCAGAATCCCAGCCAGATACTCCGC  160
C1_Allele1    GTATTTGGTCAATGGCCTCCACCCACTGACCCTGCGCTGGGAAG------------------------------------  124
C1_Allele2    GTATTTGGTCAATGGCCTCCACCCACTGA--------------------------------TCCCAGCCAGATACTCCGC  128
C2_Allele1    GTATTTGGTCAATCCCTCCCGCACAC--------------------------------------CAGCCAGATACTCCGC  122
C2_Allele2    GTATTTGGTCAATGGCCTCCACCCACTGACCCTGCGCTGGGAAGAGACCCGCACAC----------------------GC  138
C2_Allele3    --------------------------------------------------------------------------------  79

               PAM
Exo4_WT       CTGGCACCCCTCTGGTGTCCCAAGATGAGAAGAGAGATGCTGAGCTGCCGAAGAAGCGTATGCGGAAGTCAAACCCCGGC  240
C1_Allele1    --GG---TCCTCTGGTGTCCCAAGATGAGAAGAGAGATGCTGAGCTGCCGAAGAAGCGTATGCGGAAGTCAAACCCCGGC  199
C1_Allele2    CTGGCACCCCTCTGGTGTCCCAAGATGAGAAGAGAGATGCTGAGCTGCCGAAGAAGCGTATGCGGAAGTCAAACCCCGGC  208
C2_Allele1    CTGGCACCCCTCTGGTGTCCCAAGATGAGAAGAGAGATGCTGAGCTGCCGAAGAAGCGTATGCGGAAGTCAAACCCCGGC  202
C2_Allele2    CTGGCACCCCTCTGGTGTCCCAAGATGAGAAGAGAGATGCTGAGCTGCCGAAGAAGCGTATGCGGAAGTCAAACCCCGGC  212
C2_Allele3    -------------------CCAAGATGAGAAGAGAGATGCTGAGCTGCCGAAGAAGCGTATGCGGAAGTCAAACCCCGGC  140

Exo4_WT       TGGGAGAACTTGGAGAAGTTGCTAGTGTTCACCGCAGCTGGGGTGAAACCCCAGGGCAAG  300
C1_Allele1    TGGGAGAACTTGGAGAAGTTGCTAGTGTTCACCGCAGCTGGGGTGAAACCCCAGGGCAAG  259
C1_Allele2    TGGGAGAACTTGGAGAAGTTGCTAGTGTTCACCGCAGCTGGGGTGAAACCCCAGGGCAAG  268
C2_Allele1    TGGGAGAACTTGGAGAAGTTGCTAGTGTTCACCGCAGCTGGGGTGAAACCCCAGGGCAAG  262
C2_Allele2    TGGGAGAACTTGGAGAAGTTGCTAGTGTTCACCGCAGCTGGGGTGAAACCCCAGGGCAAG  278
C2_Allele3    TGGGAGAACTTGGAGAAGTTGCTAGTGTTCACCGCAGCTGGGGTGAAACCCCAGGGCAAG  200
```

B

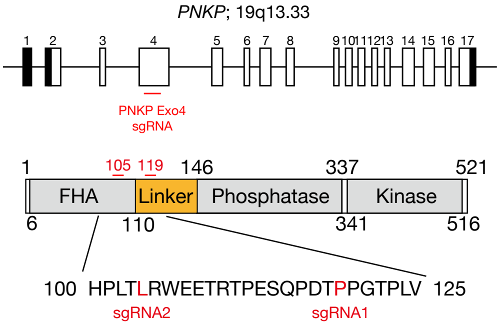
