## Supplementary figures and images for "CDKs-mediated phosphorylation of PNKP is required for end-processing of single-strand DNA gaps on Okazaki Fragments and genome stability"

### Supplementary Figure 2

Supplementary Figure 2

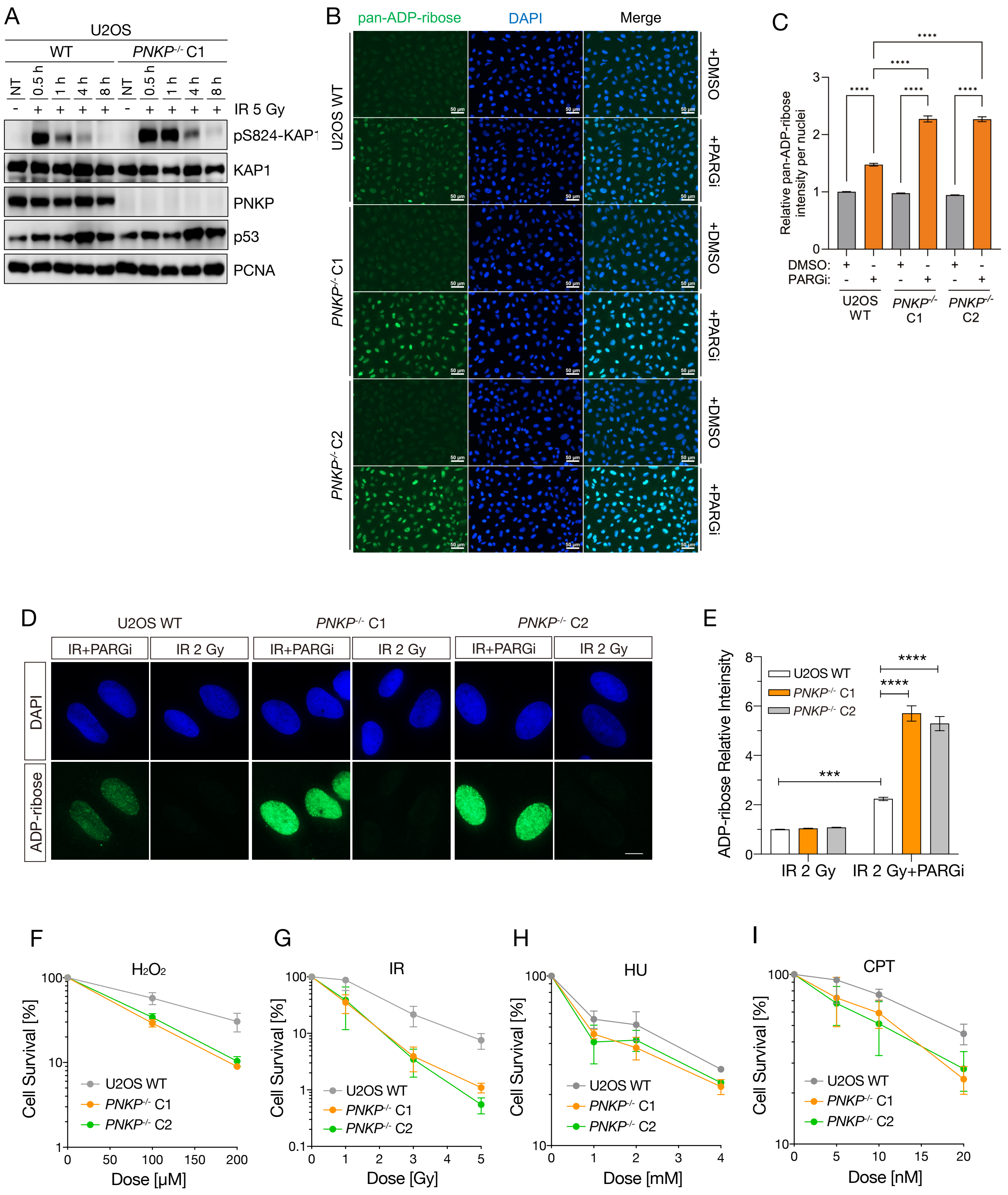

### Supplementary Figure 3

Supplementary Figure 3

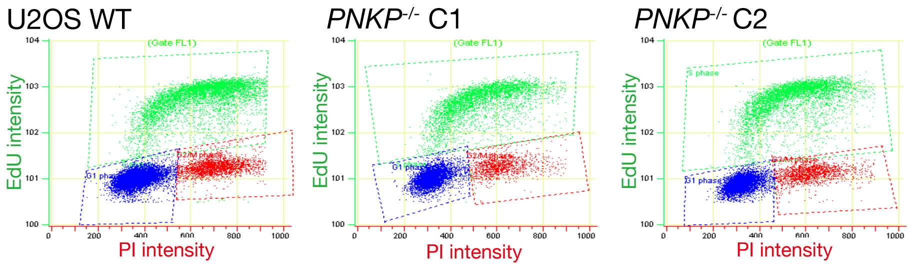

### Supplementary Figure 4

Supplementary Figure 4

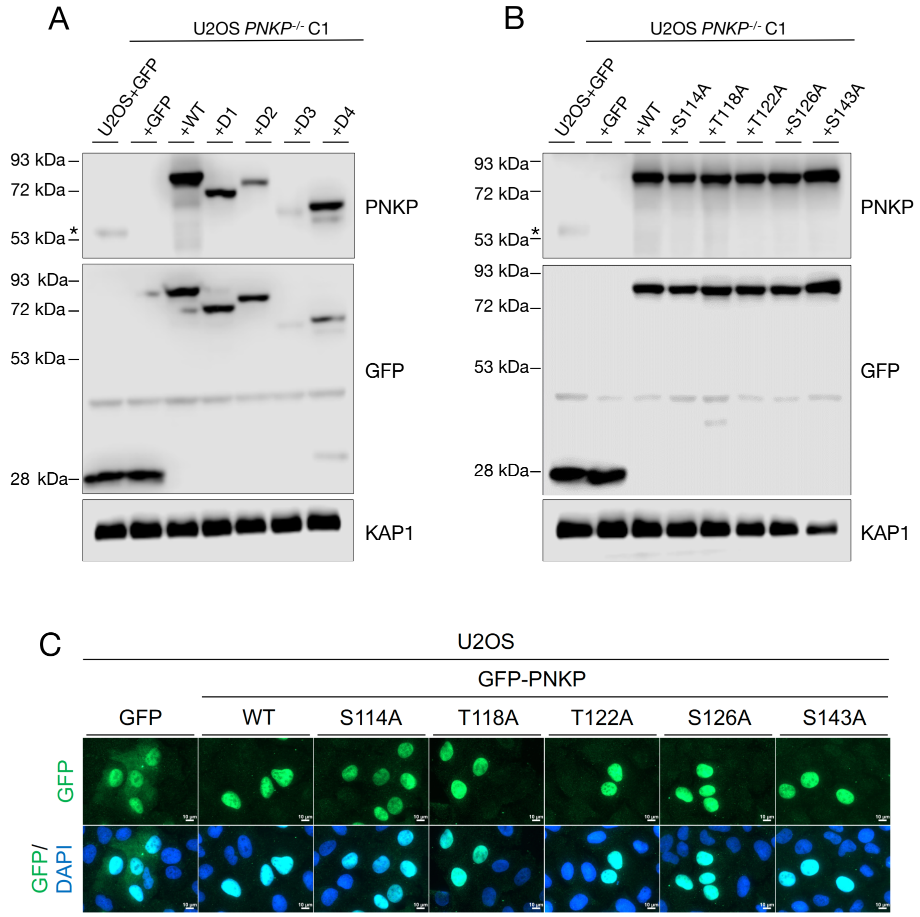

### Supplementary Figure 5

Supplementary Figure 5

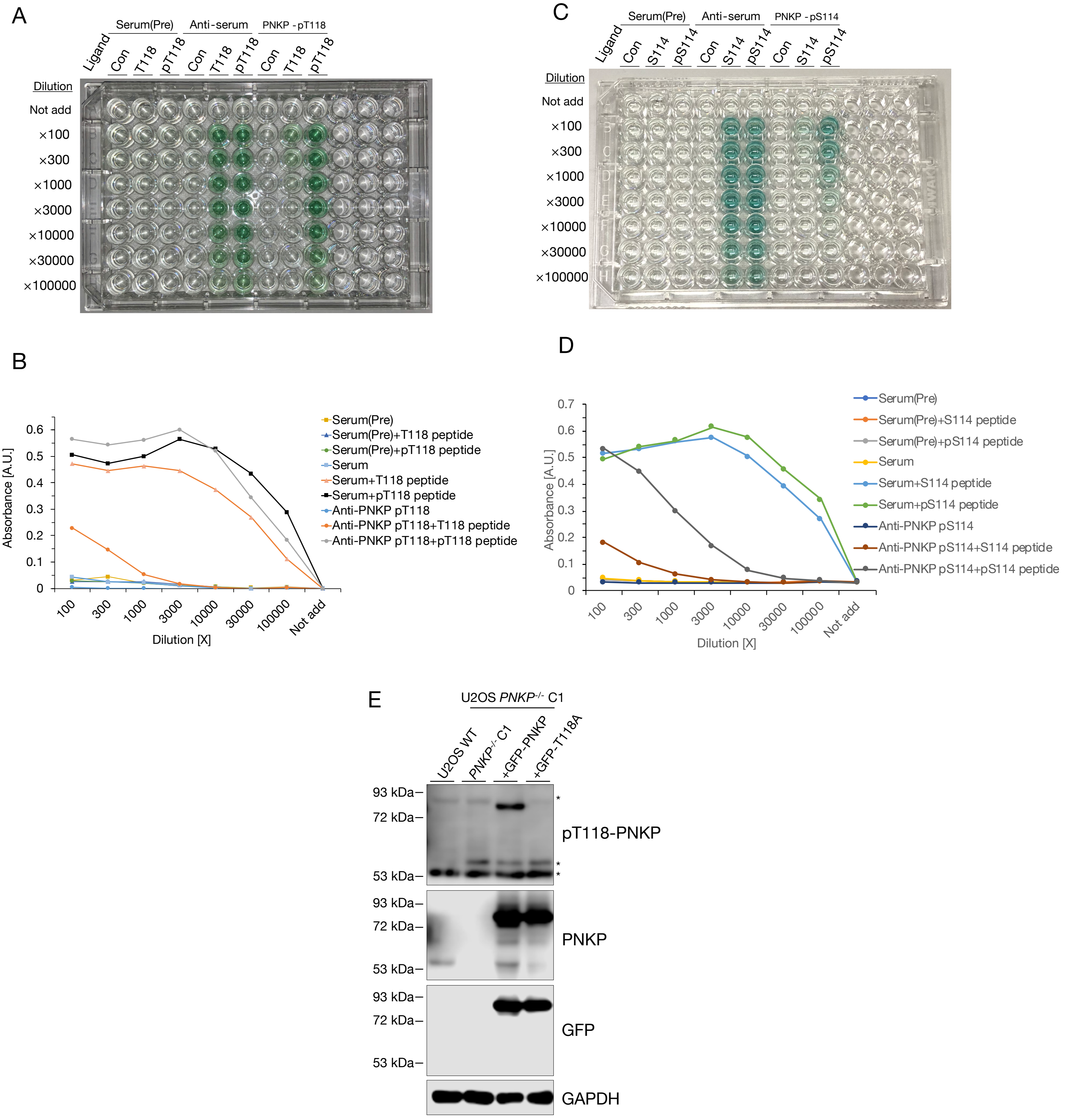

### Supplementary Figure 6

Supplementary Figure 6

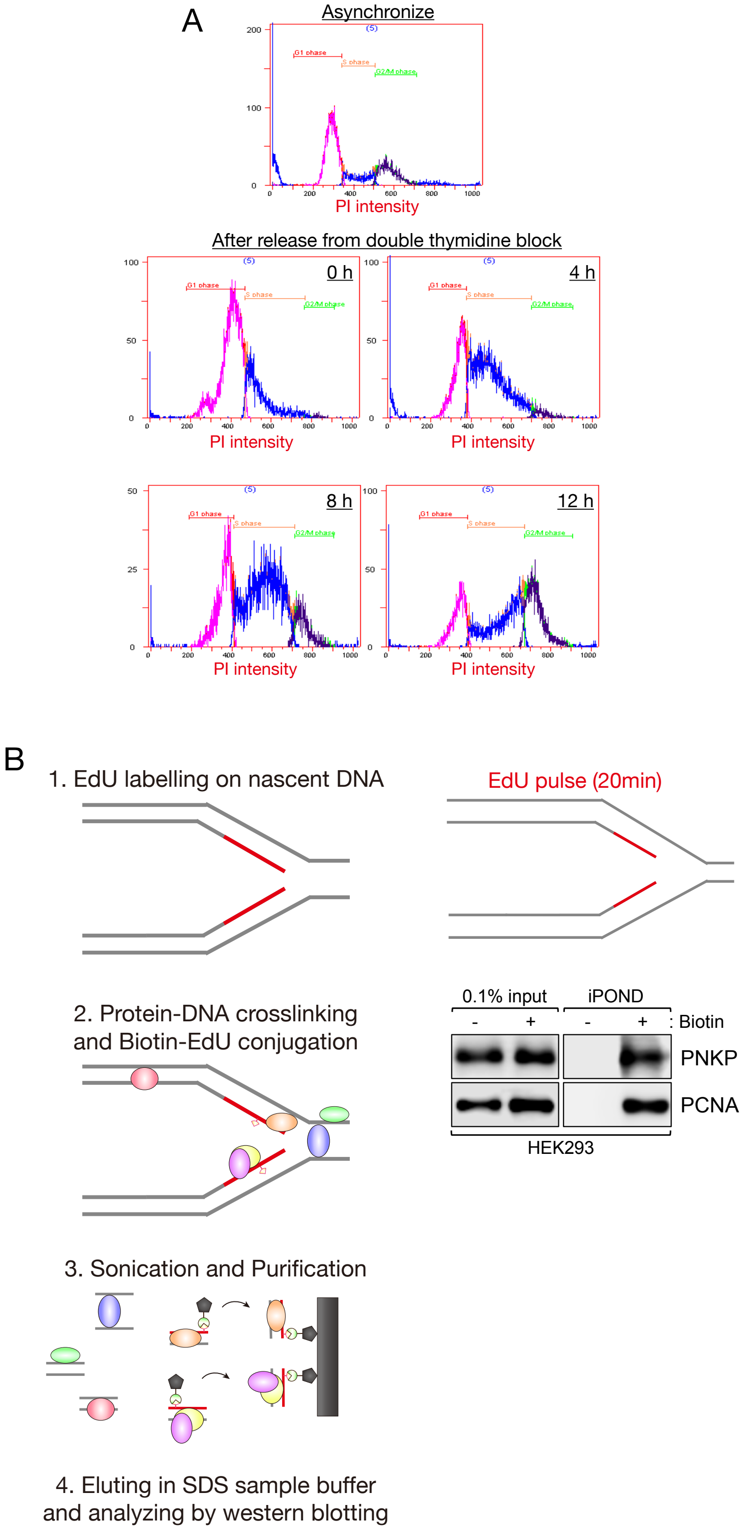

### Supplementary Figure 7

Supplementary Figure 7

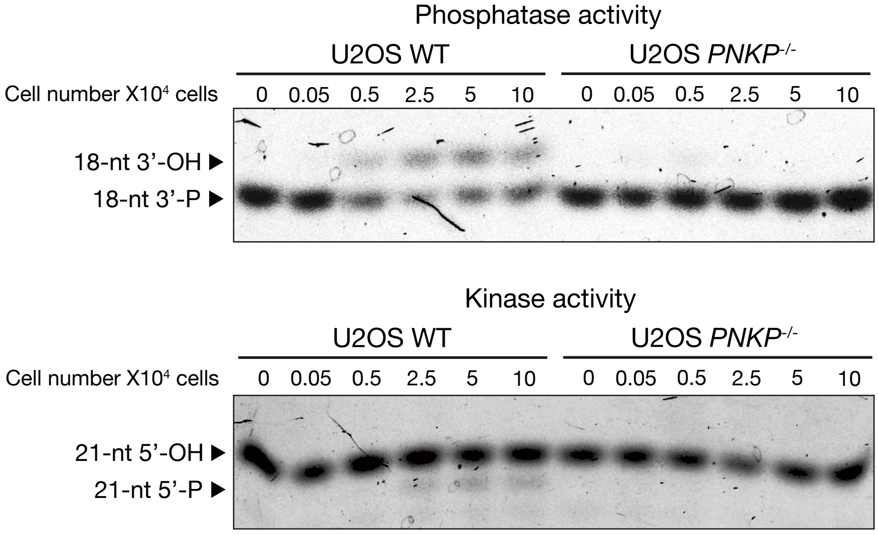

### Supplementary Figure 8

Supplementary Figure 8

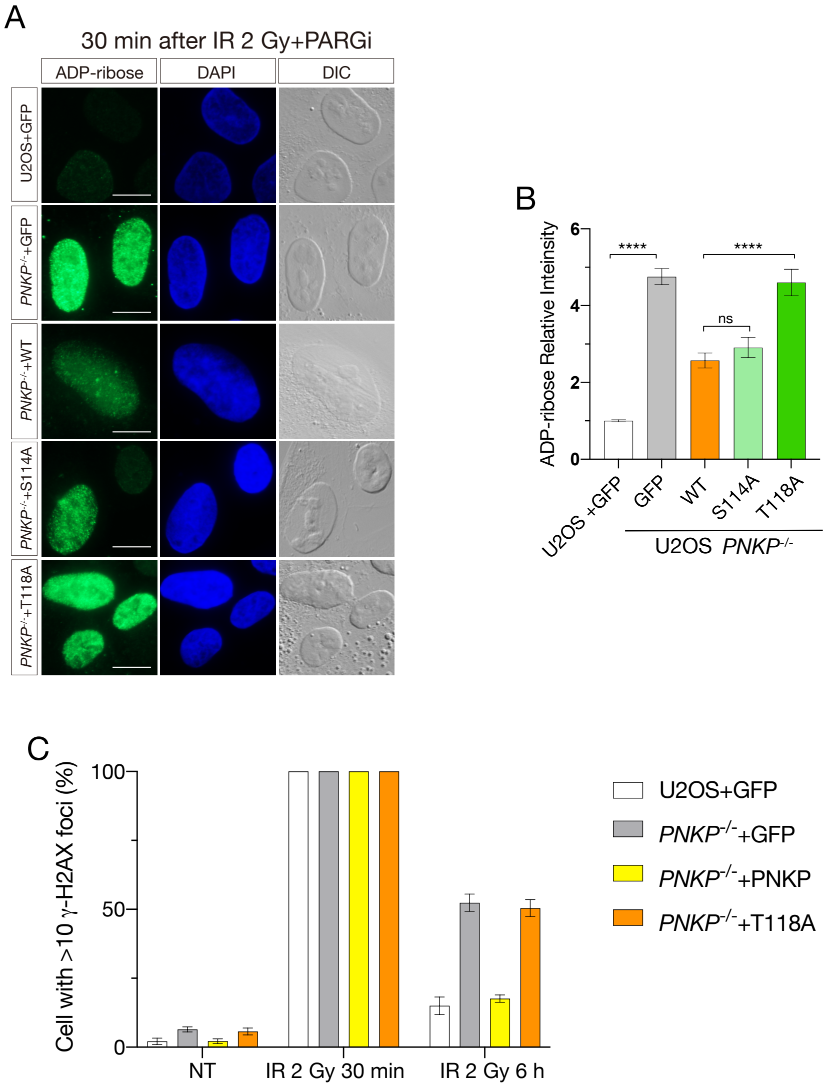

### Supplementary Figure 9

Supplementary Figure 9

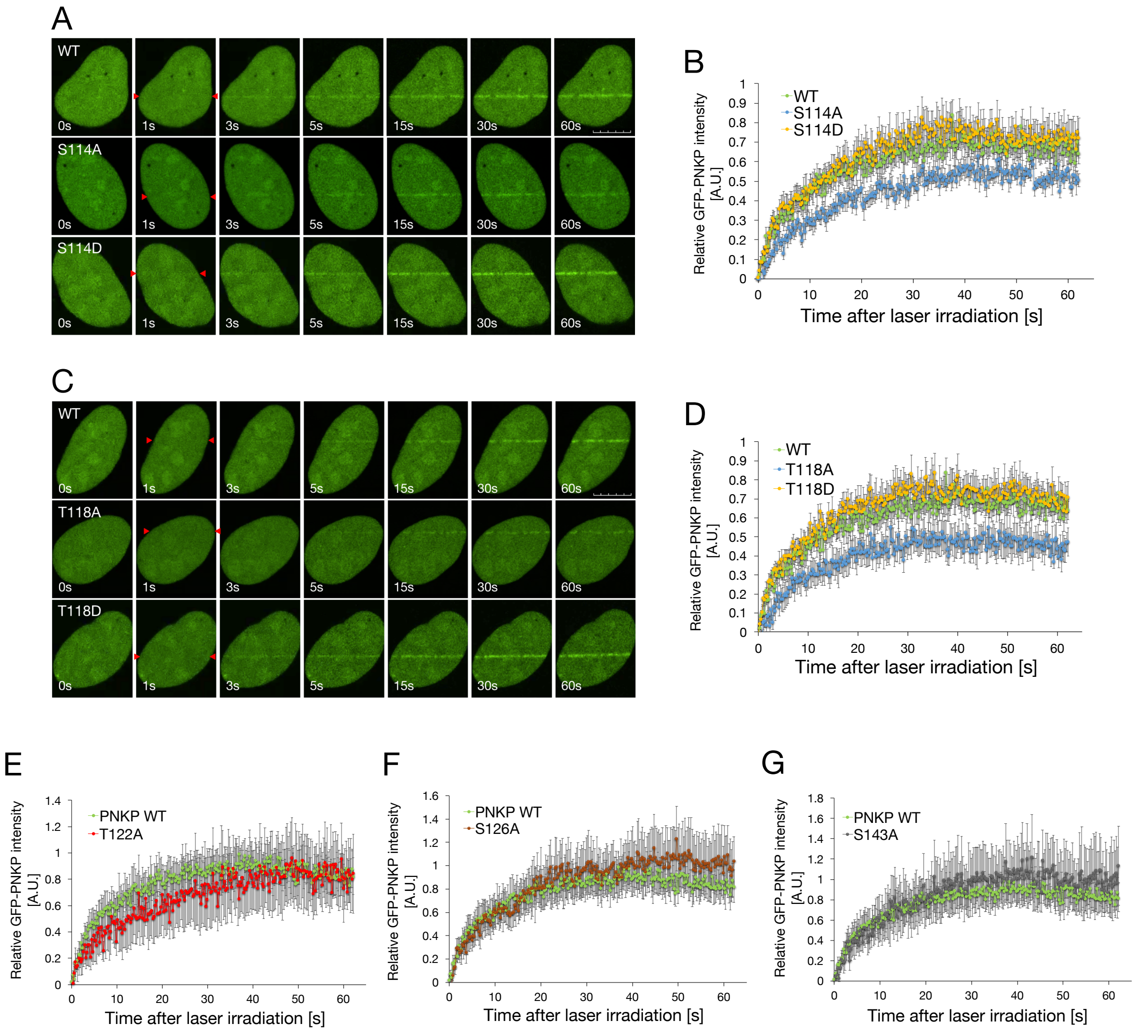
