## Supplementary Table 1 for "CDKs-mediated phosphorylation of PNKP is required for end-processing of single-strand DNA gaps on Okazaki Fragments and genome stability"

Sequences of DNA oligonucleotide primers for mutagenesis of PNKP. “F” and “R” indicate forward and reverse sequences, respectively.

| Name | | DNA sequences |
|---|---|---|
| D1 | F | CGGACTCAGATCTATGACACCAGAATCCCAGC |
| | R | GCTGGGATTCTGGTGTCATAGATCTGAGTCCG |
| D2 | F | GGGAAGAGACCCGCGGCTGGGAGAACTT |
| | R | AAGTTCTCCCAGCCGCGGGTCTCTTCCC |
| D3 | F | GCGGAAGTCAAACCCCCTCCCAGCCTTTGATC |
| | R | GATCAAAGGCTGGGAGGGGGTTTGACTTCCGC |
| D4 | F | CAGCCGGCTTCGAGTGAGTCGACGGTAC |
| | R | GTACCGTCGACTCACTCGAAGCCGGCTG |
| R35A | F | CCTGGTCCTGGGCGCGGGACCCCTGACC |
| | R | GGTCAGGGGTCCCGCGCCCAGGACCAGG |
| R48A | F | GGACCGGAAGTGCTCCGCAACTCAAGTGGAGCTG |
| | R | CAGCTCCACTTGAGTTGCGGAGCACTTCCGGTCC |
| S114A | F | GACCCGCACACCAGAAGCCCAGCCAGA |
| | R | TCTGGCTGGGCTTCTGGTGTGCGGGTC |
| S114D | F | AGACCCGCACACCAGAAGACCAGCCAGATACTCC |
| | R | GGAGTATCTGGCTGGTCTTCTGGTGTGCGGGTCT |
| T118A | F | CCAGAATCCCAGCCAGATGCTCCGCCTGG |
| | R | CCAGGCGGAGCATCTGGCTGGGATTCTGG |
| T118D | F | CACCAGAATCCCAGCCAGATGATCCGCCTGGCAC |
| | R | GTGCCAGGCGGATCATCTGGCTGGGATTCTGGTG |
| T122A | F | CTCCGCCTGGCGCCCCTCTGGTG |
| | R | CACCAGAGGGGCGCCAGGCGGAG |
| S126A | F | GCACCCCTCTGGTGGCCCAAGATGAGAAG |
| | R | CTTCTCATCTTGGGCCACCAGAGGGGTGC |
| S143A | F | GCGTATGCGGAAGGCAAACCCCGGCTG |
| | R | CAGCCGGGGTTTGCCTTCCGCATACGC |
| D171A | F | GTGGCTGGCTTTGCTCTGGACGGGACG |
| | R | CGTCCCGTCCAGAGCAAAGCCAGCCAC |
| K378A | F | CCCTGGGGCCGGGGCGTCCACCTTTCTC |
| | R | GAGAAAGGTGGACGCCCCGGCCCCAGGG |
