## Supplementary Table 2 for "CDKs-mediated phosphorylation of PNKP is required for end-processing of single-strand DNA gaps on Okazaki Fragments and genome stability"

Supplementary table 2
Oligonucleotide sequences of siRNAs for specific depletion with indicated proteins.

| Name | | Oligonucleotides | Target site (5'-3') |
|---|---|---|---|
| siLUC | Sense | | TAAGGCTATGAAGAGATAC Dharmacon, cat# D-001210-02-20 |
| | Anti-sense | | |
| siPNKP 3'-UTR Ver.1 | Sense | CACAAUAAACGCUGUUUCUCC | CACAATAAACGCTGTTTCTCC |
| | Anti-sense | AGAAACAGCGUUUAUUGUGGA | |
| siPNKP 3'-UTR Ver.2 | Sense | ACAAUAAACGCUGUUUCUCCUUG | ACAATAAACGCTGTTTCTCCTTG |
| | Anti-sense | AGGAGAAACAGCGUUUAUUGUGG | |
